# TGF-beta/Activin ligand Myoglianin couples muscle growth to the initiation of metamorphosis

**DOI:** 10.1101/2020.04.21.053835

**Authors:** Lorrie L. He, Sara Hyun Joo Shin, Zhou Wang, Isabelle Yuan, Ruthie Weschler, Allie Chiou, Takashi Koyama, H. Frederik Nijhout, Yuichiro Suzuki

**Affiliations:** Department of Biological Sciences, 106 Central St., Wellesley College, Wellesley, MA 02481; Instituto Gulbenkian de Ciência, Rua da Quinta Grande, 6, 2780-156 Oeiras, Portugal; Department of Biology, University of Copenhagen, Universitetsparken 15, 2100 Copenhagen, Denmark; Department of Biology, Duke University, Durham, NC 27708

**Author notes:** denotes equal contribution (Lorrie L. He, Sara Hyun Joo Shin).

**Keywords:** Myoglianin, threshold size, body size, muscle growth, *Manduca sexta*

## Abstract

Although the mechanisms that control growth are now well understood, the mechanism by which animals assess their body size remains one of the great puzzles in biology. The final larval instar of holometabolous insects, after which growth stops and metamorphosis begins, is specified by a threshold size. We investigated the mechanism of threshold size assessment in the tobacco hornworm, *Manduca sexta*. The threshold size was found to change depending on the amount of exposure to poor nutrient conditions whereas hypoxia treatment consistently led to a lower threshold size. Under these various conditions, the mass of the muscles plus integuments was correlated with the threshold size. Furthermore, the expression of *myoglianin* (*myo*) increased at the threshold size in both *M. sexta* and *Tribolium castaneum*. Knockdown of *myo* in *T. castaneum* led to larvae that underwent supernumerary larval molts and stayed in the larval stage permanently even after passing the threshold size. We propose that increasing levels of Myo produced by the growing tissues allow larvae to assess their body size and trigger metamorphosis at the threshold size.

## BACKGROUND

In animals that undergo determinate growth, the juvenile stage, during which growth occurs, is separated from the adult stage. The two stages are often separated by a major developmental transition, such as puberty or metamorphosis, which involves physiological, morphological and behavioral changes. Because these developmental transitions occur once the juveniles have grown to a particular body size, organisms must have evolved mechanisms to assess their body size. Although the molecular, genetic and physiological mechanisms that control growth, puberty and metamorphosis are now well-known, the mechanism by which animals assess their body size and determine when to stop growing remains a major unresolved issue in developmental biology.

Holometabolous insects, insects that undergo complete metamorphosis, exhibit determinate growth. These insects grow by undergoing several molts. At the end of the last larval stage, they stop feeding and growing, and metamorphose into the pupal and adult stages that do not grow. The size a larva attains during the last larval instar therefore defines the size of the adult insect. Previous studies have shown that the decision to stop growing and begin metamorphosis is marked by the attainment of a precise body size called the threshold size (Nijhout, 1975).

The threshold size was first identified in the tobacco hornworm, *Manduca sexta* (Nijhout, 1975). *M. sexta* typically undergo five larval instars in laboratory conditions. However, when *M. sexta* larvae are fed a low-nutrient diet, larvae grow more slowly and can undergo 1, 2 or 3 additional instars before entering metamorphosis. The threshold checkpoint occurs at the beginning of each instar (Kingsolver, 2007; Nijhout, 1975). A larva below the threshold size at the beginning of an instar will undergo additional molts and increase its body size. Once a larva is above the threshold size when it molts, it enters the last larval instar and will metamorphose at the end of that instar (Kingsolver, 2007; Nijhout, 1975).

Artificial selection on body size has demonstrated that threshold size can evolve, indicating that there is a genetic component to the mechanism that determines the threshold size (Grunert et al., 2015). There are two equivalent measures that can detect threshold size: It can be measured as the width of the head capsule, or as the mass of a larva at the beginning of the instar (Grunert et al., 2015; Nijhout, 1975). Both measures are equivalent. Thus, some mechanism of size sensing must exist that is somehow associated with these measures of body size.

Recent studies have identified several molecular regulators whose disruption causes supernumerary molts or precocious metamorphosis. These molecular regulators affect the production of, or sensitivity to, juvenile hormone (JH). JH modifies the actions of the molting hormone, 20-hydroxyecdysone, to prevent the organism from progressing from one life history stage to the next (Riddiford, 1996). JH acts by binding to the basic helix-loop-helix-Per-Arnt-Sim domain protein receptor, Methoprene-tolerant (Met) (Konopova and Jindra, 2007). In flour beetle *Tribolium castaneum,* knockdown of *Met* causes larvae to have reduced sensitivity to JH and undergo early metamorphosis, forming miniature adults (Konopova and Jindra, 2007). Silencing the expression of the JH-response gene, *Krüppel homolog* (*Kr-h1*), also leads to precocious metamorphosis in this species (Minakuchi et al., 2009).

It has long been known that removing the corpora allata, the glands that secrete JH, can cause premature metamorphosis, resulting in a dwarf adult. JH-deficient larvae of holometabolous insects, including *M. sexta*, *T. castaneum* and the silkworm *Bombyx mori*, undergo precocious metamorphosis (Daimon et al., 2012; Minakuchi et al., 2008; Ohtaki et al., 1971; Suzuki et al., 2013; Tan et al., 2005). In *T. castaneum*, knockdown of *ventral veins lacking* (*vvl*) leads to precocious metamorphosis by suppressing the production of JH (Cheng et al., 2014). Recently, the ecdysone response gene, *E93*, has been shown to be necessary to terminate JH secretion in order to initiate the onset of metamorphosis; knockdown of *E93* leads to the induction of supernumerary molts in *T. castaneum* (Chafino et al., 2019). Chafino et al (2019) demonstrated that in this species, starvation before day 1 of the final instar can induce supernumerary molts whereas starvation after day 1 does not lead to supernumerary molts. Chafino et al (2019) suggested that the mass on the first day of the final instar likely corresponds to the threshold size and demonstrated that it is associated with an increase in *E93* expression (Chafino et al., 2019). While the methodology used to determine the threshold size is different from that used in *M. sexta* and the identified stage may be more consistent with the attainment of irreversible pupal commitment, it is clear that the decision to become a pupa (and hence the threshold size) is already reached 24 hrs after the molt to the final instar. Although the clearance of JH is a prerequisite for the decision to metamorphose, removal of JH is likely only to be the proximate mechanism that allows larvae to initiate metamorphosis and not part of the mechanism by which a larva *assesses* its size.

Hemimetabolous insects are characterized by incomplete metamorphosis, in which nymphs metamorphose directly into adults. To our knowledge, no study has demonstrated the existence of a threshold size in this group of insects. In these insects, JH also plays a role in the timing of adult development. Knockdown of *Kr-h1* in the penultimate nymphal instar of the firebug, *Pyrrhocoris apterus*, the German cockroach, *Blattella germanica*, and the brown planthopper, *Nilaparvata lugens*, leads to precocious adult development (Konopova et al., 2011; Li et al., 2018; Lozano and Belles, 2011). In addition, silencing *myoglianin* (*myo*), a gene coding for one of the ligands of the TGF-beta/Activin signaling pathway, has been shown to cause extra nymphal molts (Ishimaru et al., 2016; Kamsoi and Belles, 2019): Increased expression of *myo* has been shown to trigger the final nymphal instar in *Blattella germanica* (Kamsoi and Belles, 2019). Myo expression in the corpora allata/corpora cardiaca in the fifth (penultimate nymphal) instar nymphs leads to the repression of the JH biosynthesis gene, *jhamt,* in the sixth (last) instar nymphs (Kamsoi and Belles, 2019). Likewise, in the field cricket, *Gryllus bimaculatus,* RNA interference (RNAi)-mediated knockdown of *myo* leads to supernumerary molts accompanied by an increase in *jhamt* expression (Ishimaru et al., 2016). These studies demonstrate that Myo regulates JH production in hemimetabolous insects. In addition, Myo has been implicated in ecdysteroid production as the expression of the ecdysone biosynthesis gene *neverland* decreases in response to *myo* RNAi (Kamsoi and Belles, 2019).

The role of *myo* in holometabolous insects is not well known except for its functions in muscles (Augustin et al., 2017; Lo and Frasch, 1999). Myo shares a 46% amino acid sequence identity with the vertebrate Bone Morphogenetic Protein 11 (BMP11 or Growth differentiation factor 11 (GDF11)) and Myostatin (or GDF8) (Lo and Frasch, 1999). In vertebrates, Myostatin was first isolated in mice and characterized for its role in halting skeletal muscle growth (McPherron et al., 1997; Whittemore et al., 2003). In *Drosophila,* Myo is expressed in embryonic muscles as well as glial cells (Lo and Frasch, 1999). During the larval stage, Myo acts like Myostatin at the neuromuscular junction (NMJ) to suppress synaptic transmissions and NMJ growth and branching (Augustin et al., 2017). In addition, knockdown of muscle-derived *myo* leads to increased muscle size and overall body size, indicating that Myo inhibits muscle growth (Augustin et al., 2017). However, *myo* mutant *Drosophila* larvae do not undergo extra larval molts (Augustin et al., 2017). In fact, unlike most other insects, the final instar *Drosophila* larvae do not respond readily to JH; topical application of JH fails to dramatically delay metamorphosis and does not induce supernumerary molts (Riddiford and Ashburner, 1991). Thus, identification of threshold size is challenging in this species. In summary, although molecular disruptions that cause precocious metamorphosis or supernumerary molts have been identified, none of these answer the question of how size is assessed so that metamorphosis occurs at the correct species-specific body size.

In this study, we sought to examine how *M. sexta* larvae assess their body size to initiate metamorphosis. Our approach was to utilize two distinct methods to generate a wide range of body sizes at the end of the fourth instar (penultimate instar under laboratory conditions): nutrient-deprivation and hypoxia. As mentioned above, nutrient-deprivation is the standard way by which threshold size has been determined in *M. sexta*. Aside from nutrient deprivation, previous studies have shown that hypoxia can also stunt growth in most insect species, including *M. sexta* (Callier and Nijhout, 2011; Frazier et al., 2001; Greenberg and Ar, 1996). We therefore exposed larvae at different instars to hypoxic conditions to generate a range of fourth instar larval sizes. We then sought to identify traits that correlate with the attainment of the threshold size under both nutrient-deprivation and hypoxia conditions to narrow down the potential molecular regulators involved in threshold size determination.

We found that nutrient deprivation at different times of larval development leads to distinct threshold sizes and found that the relative muscle/integument mass is correlated with the attainment of the threshold size. We also show that *myo* is expressed differentially in muscles/integuments of pre-threshold size larvae relative to that of post-threshold size larvae. Because knockdown of gene expression *in vivo* is not possible in *M. sexta*, we explored the function of *myo* in another holometabolous insect, the flour beetle *T. castaneum*, where RNAi is possible (Tomoyasu and Denell, 2004). We demonstrate that *myo* dsRNA-injected larvae continue to molt into supernumerary larval instars and never initiate metamorphosis even after surpassing the threshold size, indicating that *myo* is the signal by which body size is sensed and that triggers the transition between growth and metamorphosis.

## RESULTS

### Effect of nutrient deprivation on the threshold size

Under standard laboratory rearing conditions, *M. sexta* larvae invariably undergo five larval instars. This precludes us from determining the threshold size. Thus, the standard way to determine the threshold size is to temporarily starve larvae or to feed larvae a diet containing a reduced amount of protein to generate large variability in size (Grunert et al., 2015; Nijhout, 1975). Here, we used both methods to determine the threshold size. First, larvae were fed an experimental diet with reduced levels of protein (40% diet) starting the second instar. These larvae had a threshold size of approximately 0.85 g (Fig. 1A, 1C). In contrast, shifting the timing of nutrient deprivation altered the threshold size. When larvae were starved during the third instar after one day of feeding, or when third instar larvae were placed on 40% diet temporarily and returned to the normal diet at the end of the third instar, larvae had a reduced threshold size: Larvae on the 40% diet only in the third instar and larvae starved in the third instar had a threshold size of approximately 0.65 g and 0.75 g, respectively (Fig. 1A, C). Thus, the timing of nutrient deprivation appears to affect the threshold size.

**Figure 1.**
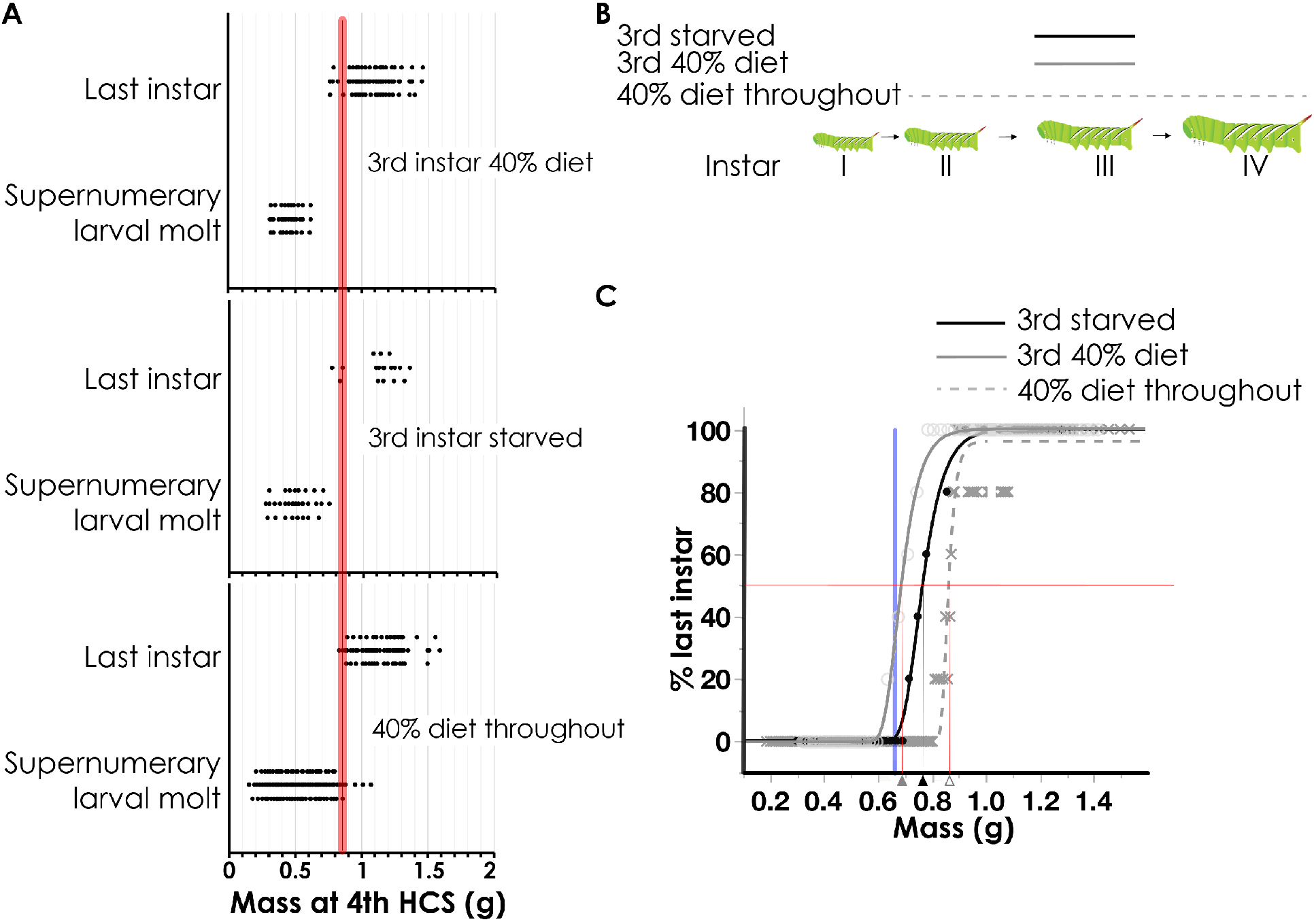
Nutritional conditions affect threshold size. (A) A plot of mass at the end of the fourth instar vs developmental fate. (Top) Larvae fed a 40% diet during the third instar. (Middle) Larvae starved during the third instar starting day 1. (Bottom) Larvae fed a 40% diet throughout the larval instar. Red line indicates the threshold size for larvae fed a 40% diet throughout much of their larval life. “Supernumerary larval molt” indicates that the fifth instar larvae molted into another larval instar. “Last instar” indicates that the larvae initiated wandering at the end of the fifth instar. Larvae were tracked until they exhibted signs of pupation or initiated a supernumerary molt. (B) Experimental scheme. (C) The average masses of five larvae at the end of the fourth instar were plotted against the percentage of larvae that entered final larval instar. Solid black line represents larvae that were starved during the third instar. Solid gray line represents larvae that were fed a 40% diet during the third instar. Dotted line represents larvae that were fed a 40% diet during the majority of the growth period. Triangles indicate the mass where 50% of larvae entered the final instar (i.e. threshold size). Lines represent Gompertz 3P model fits. Blue line represents threshold size of hypoxia-treated larvae.

### Hypoxia generates supernumerary larvae

Hypoxia has also been demonstrated to slow down the growth rate (Callier et al., 2013; Harrison et al., 2015). Therefore, we exposed third instar larvae to hypoxia to see if we could generate smaller larvae. After exposure to hypoxia, fourth instar larvae were returned to normoxic conditions and weighed daily until the beginning of the wandering stage, which indicates the onset of metamorphosis (Fig. S1A). These larvae grew to a smaller size at the end of the fourth instar and had two distinct fates: at the end of the fifth instar, 40% of the larvae (n=61) underwent a supernumerary larval molt, and the remaining larvae entered the wandering stage (n=90) (Fig. S1B). Larvae exposed to hypoxic conditions during the third instar that wandered at the end of the fifth instar had a fourth instar feeding period of 3.0 days, similar to the average feeding time of normoxic control larvae, which took 2.9 days (Fig. S2). In contrast, larvae that underwent a supernumerary molt at the end of the fifth instar had a significantly reduced fourth instar feeding period of 2.0 days (Fig. S2; One-way ANOVA: F(2,159)=125.556, p<0.0001). Thus, the feeding duration of the fourth instar is a significant predictor of the nature of the molt that occurs at the end of the fourth instar.

The weights at the end of the fourth instar are highly predictive of the fate of the molt, indicating that the threshold size checkpoint occurs at the end of the fourth instar, similar to the nutrient-deprivation conditions (Fig. S3A, B). The mass at the end of the third instar is not predictive of the mass at the end of the fourth instar (Fig. S3C). In the hypoxia-treated larvae, the threshold size is approximately 0.65 g (Fig. 2, S3B, C), similar to that of larvae reared under normoxia/low-nutrient conditions only during the third instar.

**Figure 2.**
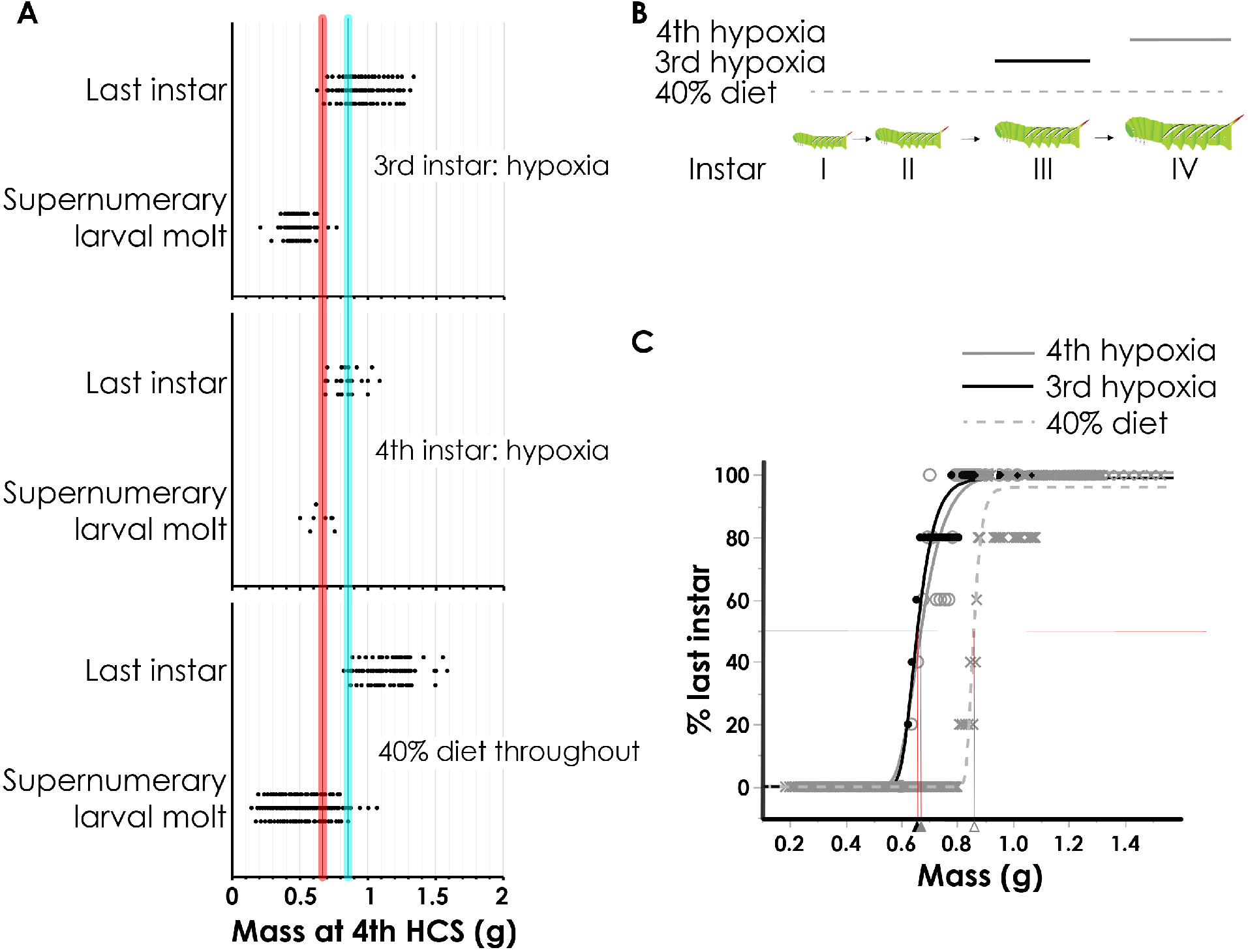
Third and fourth instar hypoxia-treated larvae have similar threshold sizes whereas 40% diet delays the attainment of threshold size. (A) A plot of mass at the end of the fourth instar vs developmental fate. Red line indicates the threshold size for the hypoxia-treated larvae. Blue line indicates the threshold size for the 40% diet fed larvae. “Supernumerary larval molt” indicates that the fifth instar larvae molt into another larval instar. “Last instar” indicates that the larvae initiated wandering at the end of the fifth instar. (B) Scheme of experimental treatments. Lines represent duration of treatments. Larvae were subjected to hypoxic conditions for the duration of the third or fourth instar, or fed a 40% diet starting in early larval life, and tracked until they exhibited signs of wandering or initiated a supernumerary molt. (C) The moving average masses of five larvae at the end of the fourth instar were plotted against the percentage of larvae that entered final larval instar. Solid black line represents larvae that were placed in hypoxic condition during the third instar. Solid gray line represents larvae that were placed in hypoxic condition during the fourth instar. Dotted line represents larvae that were fed a 40% diet during the majority of the growth period. Triangles indicate the mass where 50% of larvae entered the final instar (i.e. the threshold size). Lines represent Gompertz 3P model fits.

Since our nutrient deprivation study indicated that the threshold size can shift depending on the timing of nutrient deprivation, we reared fourth instar larvae in hypoxic conditions instead of third instar larvae and observed whether the threshold size changed. Under these conditions, the threshold size was also approximately 0.65 g, similar to the threshold size of larvae exposed to hypoxia during the third instar (Fig. 2A, C). Thus, it appears that the timing of hypoxia treatment does not alter the threshold size.

### Muscle mass is correlated with threshold size attainment

Because we determined that distinct nutritional conditions can affect the threshold size, we hypothesized that the size of a growing body part might contribute to threshold size determination. Two tissues that grow extensively throughout an instar are the fat body and the muscles. We therefore determined the muscle/integument mass and fat body mass for the third instar hypoxia-treated larvae and larvae fed a 40% diet throughout much of the early larval instars as these two treatments reliably generate fourth instar larvae with masses close to the threshold size and lead to distinct threshold sizes. We also determined the muscle/integument mass and fat body mass at the end of the third and fourth instar larvae reared under normoxia/normal diet conditions.

Although the third and fourth instar normoxia/normal diet-fed larvae are at the extreme ends of the size range, the hypoxia-treated and normoxia/normal diet-fed larvae appear to have similar muscle/integument mass relative to the wet mass, indicating that the muscle grows similarly in hypoxia-treated and normoxia/normal diet-fed larvae (Fig. 3A). In contrast, the larvae fed a 40% diet had smaller relative mass of the muscles/integuments than hypoxia-treated and normoxia/normal diet-fed larvae (Fig. 3A). We found that the relative mass of the muscles/integuments was larger for hypoxia-treated larvae than larvae fed a 40% diet, and a significant wet mass X treatment interaction was observed (ANCOVA: F(1,39) = 13.016, p<0.001; Fig. 3A). Intriguingly, the mass of the muscles/integuments at the threshold size for both the hypoxia-treated larvae and the 40% diet-fed larvae were similar. In contrast, fat body mass scaled with whole body mass similarly in both environmental conditions, and there was no significant wet mass X treatment interaction (ANCOVA: F(1,39) = 0.0142, p = 0.906; Fig. 3B). Together, these results suggest that muscle/integument mass is correlated with attainment of threshold size.

**Figure 3.**
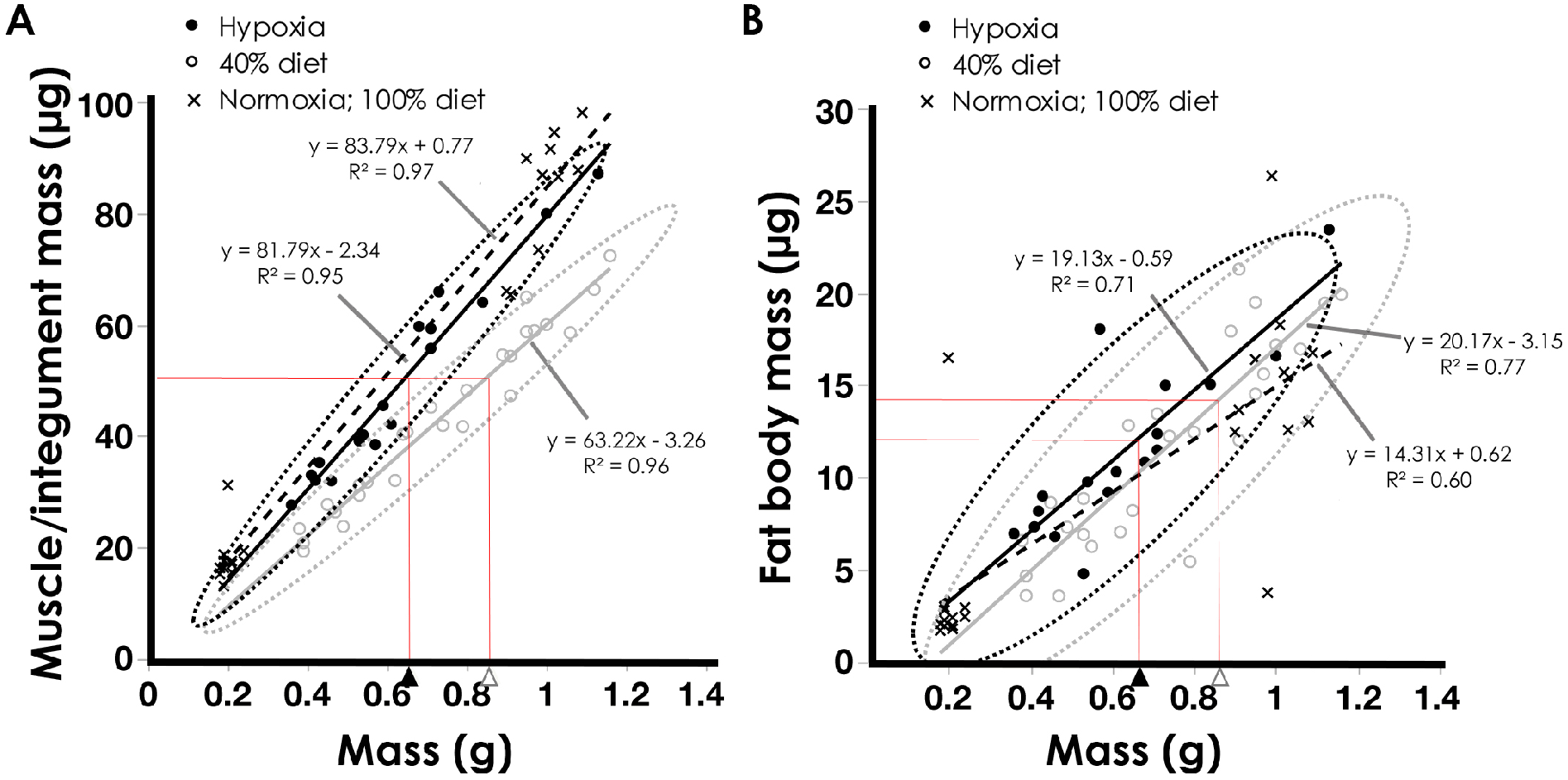
The relative dry mass of muscles/integuments is reduced in larvae fed a 40% diet relative to hypoxia-treated larvae. (A) Dry mass of muscles/integuments plotted against wet mass of fourth instar larvae. (B) Dry mass of fat body plotted against wet mass of fourth instar larvae. Red lines represent the relationship between the threshold sizes (triangles) and muscle/integument mass or fat body mass at the threshold sizes. Larvae were either placed in hypoxic conditions during the third instar or reared on a 40% diet throughout much of the larval stage. All dissections were performed at the end of the fourth instar. The dotted confidence ellipses are drawn at the 95% confidence level.

### *myo* expression in muscles is correlated with the attainment of threshold size

Given that the integument/muscle mass was correlated with the attainment of threshold size, we hypothesized that the growing muscles might produce a factor that signals the swtich from the growth phase to the metamorphic onset. We reasoned that this signal would have to be expressed differentially between pre-threshold size and post-threshold size larvae reared under normoxia/normal diet, hypoxia and nutrient-deprivation conditions. We also suspected that the signal might be secreted in order to be able to communicate with the rest of the body. Finally, we reasoned that this factor would need to be expressed in the muscle. Given that Myo is expressed in the muscles (Augustin et al., 2017; Lo and Frasch, 1999) and its removal leads to supernumerary molts in hemimetabolous insects (Kamsoi and Belles, 2019), we suspected that Myo might be a potential candidate factor. We identified three Activin ligand genes in the *M. sexta* genome. The predicted amino acid sequences were used to verify the identity of the Myo-coding gene (Fig. S4). The alignment of *M. sexta* Myo is shown in Fig. S5.

We first examined the *myo* expression in the anterior portion (containing the brain and thoracic structures) of second, third and fourth instar larvae undergoing a molt using quantitative RT-PCR (qPCR). We found that the expression of *myo* increased with each molt (Fig. 4A). To examine how the tissue-specific expression of *myo* changes between larvae at the end of the third instar (under the threshold size) and those at the end of the fourth instar (above the threshold size), the muscles, fat body and CNS were dissected from larvae reared under standard rearing conditions. We observed a significant increase in the expression of *myo* in the muscles (Student’s t-test: t(6)=2.82, p<0.05), but no significant increase was detected in the fat body (Student’s t-test: t(6)=0.637, p=0.55) and the CNS (Student’s t-test: t(5)=0.417, p=0.69) (Fig. 4B-D). Thus, an increase in *myo* expression in the muscle was correlated with the attainment of threshold size in normoxia/normal diet-fed larvae.

**Figure 4.**
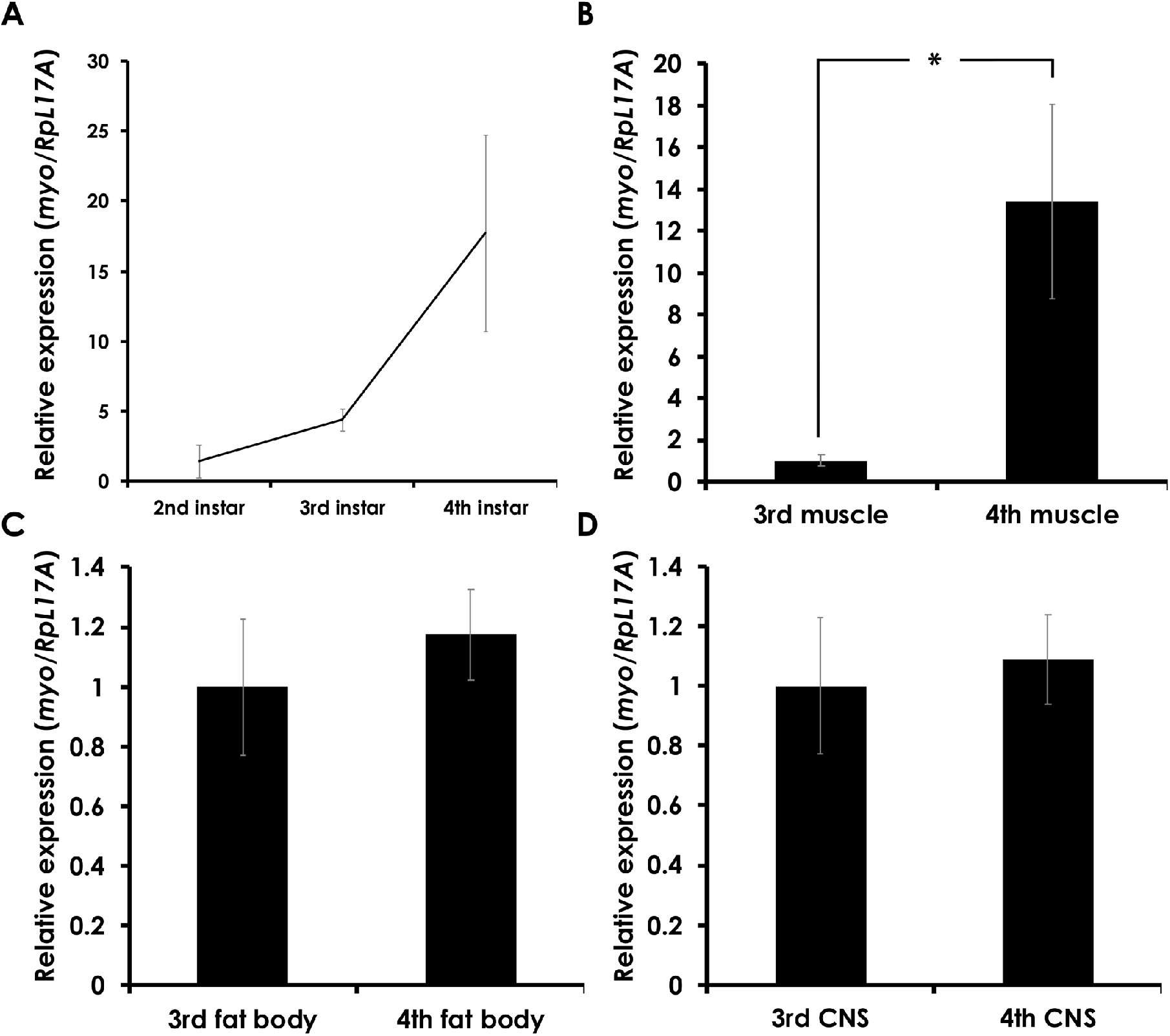
Expression of *myo* in normoxia/normal diet-fed *M. sexta* larvae. (A) *Myo* expression in the anterior half of the whole body was determined at HCS of the second, third and fourth instar of untreated larvae. Error bars represent standard error. Expression represents mean of three biological replicates, each with three technical replicates. (B-D) *Myo* expression in the muscles (B), fat body (C) and CNS (D) of normoxia/normal diet-fed third and fourth instar larvae. Error bars represent standard error. Expression represents mean of three or four biological replicates, each with three technical replicates. * indicates a statistically significant difference (Student’s t-test: p<0.05).

To further explore this correlation, we examined the tissue-specific expression of *myo* in muscles, fat body and CNS of the pre- and post-threshold size larvae in both hypoxia-treated and 40% diet-fed fourth instar larvae. We found significantly higher expression of *myo* in muscles of post-threshold size larvae than in pre-threshold size larvae in both hypoxia-treated and 40% diet fed larvae (Fig. 5; Student’s t-test: t(8)=3.65, p<0.01 for hypoxia; t(8)=3.98, p<0.005 for 40% diet). In the fat body, significantly higher expression of *myo* was observed in the post-threshold size fat body of 40% diet fed larvae relative to pre-threshold size larvae (Student’s t-test: t(7)=3.41, p<0.05). However, in hypoxia-treated larvae, no statistically significant differences were observed although post-threshold size larvae tended to have higher *myo* expression (Student’s t-test: t(8)=1.43, p=0.19). In contrast, no difference in *myo* expression was observed in the CNS of pre- or post-threshold size larvae under both hypoxia and 40% diet conditions (Fig. 5; Student’s t-test: t(8)=1.01, p=0.34 for hypoxia; t(8)=1.09, p=0.31 for 40% diet).

**Figure 5.**
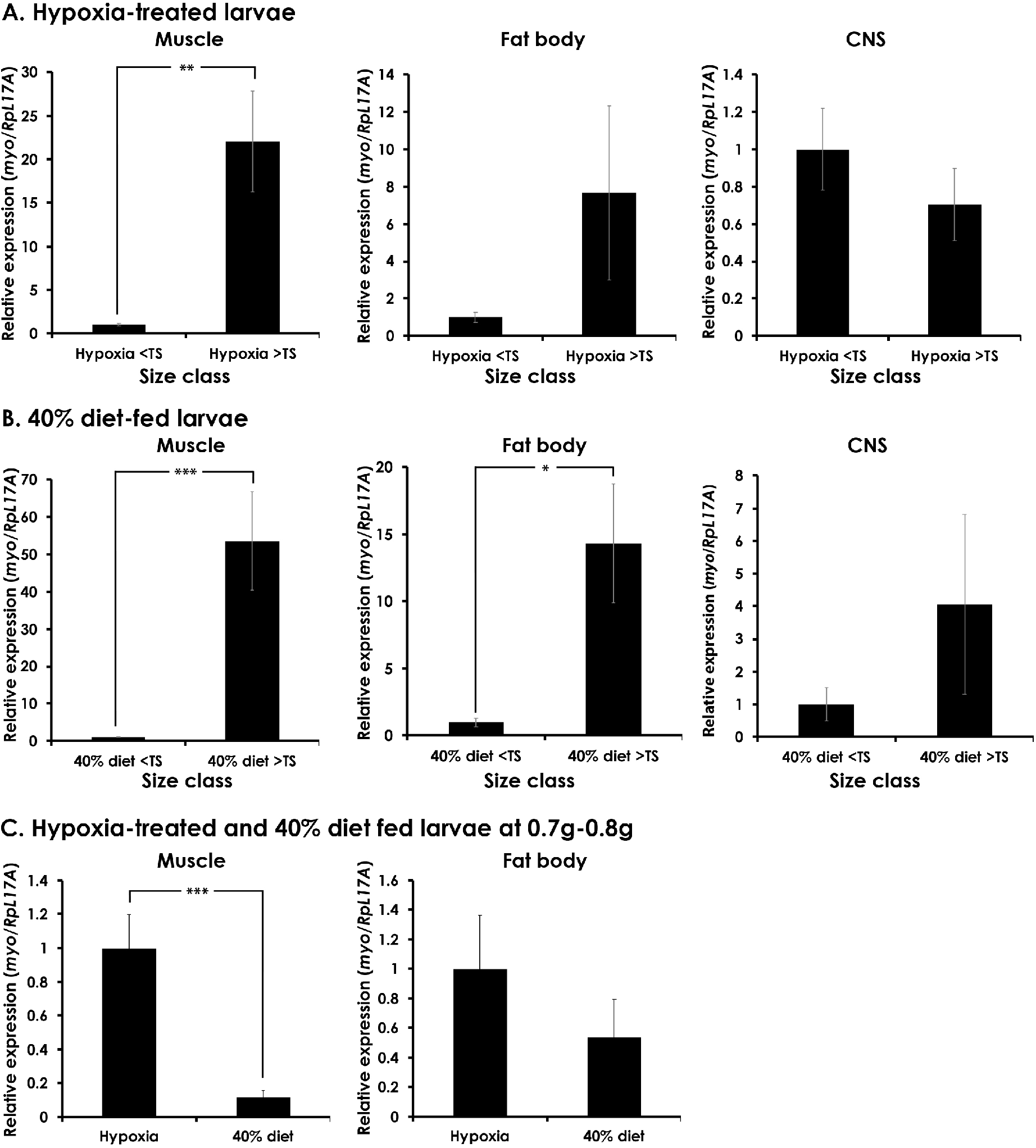
*myo* expression in muscles is signficantly elevated in post-threshold size larvae of both hypoxia-treated larvae and larvae fed a 40% diet. (A) *myo* expression in muscles, fat body and the CNS of hypoxia-treated larvae. Larvae were reared under hypoxa conditions during the third instar. (B) *myo* expression in muscles, fat body and the CNS of larvae fed a 40% diet. (C) *myo* expression in muscles and fat body of hypoxia-treated and 40% diet-fed larvae weighing 0.7 g to 0.8 g. At this weight range, hypoxia-treated larvae are above the threshold size whereas 40% diet-fed larvae are below the threshold size. For all samples, larvae were dissected at the end of the fourth instar when the larvae began to initiate a molt. Expression represents mean of 4-5 biological replicates. Each biological replicate was run with three technical replicates. Student’s t-test: * denotes p<0.05; ** denotes p<0.01; *** denotes p<0.005.

To confirm the correlation between *myo* in the muscles and the atainment of threshold size, we isolated muscles and fat body from hypoxia-treated and 40% diet-fed larvae weighing between 0.7g and 0.8g. In this weight range, hypoxia-treated larvae are above the threshold size whereas 40% diet-fed larvae are below the threshold size. We found that *myo* expression was significantly higher in the muscles of hypoxia-treated larvae than that of 40% diet-fed larvae (Fig. 5C; Student’s t-test: t(6)=4.36, p<0.005). In contrast, *myo* expression in the fat body did not differ significantly (Fig. 5C; t(6)=1.05, p=0.34). Taken together, the attainment of threshold size is consistently correlated with elevated *myo* expression in the muscles/integuments under all experimental conditions.

### JH does not shift the threshold size

Previous studies have suggested that a decline in JH may underlie threshold size attainment in other insects (Chafino et al., 2019). To determine whether JH affects threshold size attainment in *M. sexta*, larvae were placed in hypoxic conditions during the third instar until HCS and then treated with 10 *μ*g methoprene, a JH analog, two days later. These larvae typically underwent a molt approximately 1-3 days after treatment. The fates of the larvae were tracked to see if they underwent an extra molt or initiated wandering. We found that the threshold size was around 0.67 g (Fig. S6A). This is similar to that of the hypoxia-treated larvae without methoprene treatment, indicating that methoprene treatment does not shift the threshold size. All methoprene-treated larvae, including those that were destined to undergo metamorphosis at the end of the fifth instar, showed complete loss of melanic markings that are normally visible in the fifth instar (Fig. S6B, C), suggesting that JH signaling was active. Thus, while methoprene clearly had the expected effect on the body coloration, it did not alter the threshold size. Together, these data indicate that *myo* expression is correlated with the attainment of threshold size, and that methoprene treatment before threshold size attainment is insufficient to shift the threshold size.

### *myo* knockdown in *Tribolium* leads to indefinite molts

Because gene manipulation is not possible in *M. sexta*, we explored the role of *myo* in *T. castaneum,* a species where RNAi is possible. In *Tribolium*, even under normal rearing conditions, the total number of instars can vary, and like in *M. sexta,* the timing of metamorphosis is determined by a drop in JH. In our laboratory, the GA-1 strain typically undergoes 7 or 8 instars. A recent study has shown that by day 1 of the final instar, the decision to pupate and hence the threshold size has already been reached (14). We weighed larvae 1 day after a molt and determined their fates. When larvae weighed above 2.1 g, 50% of the larvae initiated metamorphosis (Fig. 6A), indicating that if larvae are above 2.1 g, they have already reached the threshold size.

**Figure 6.**
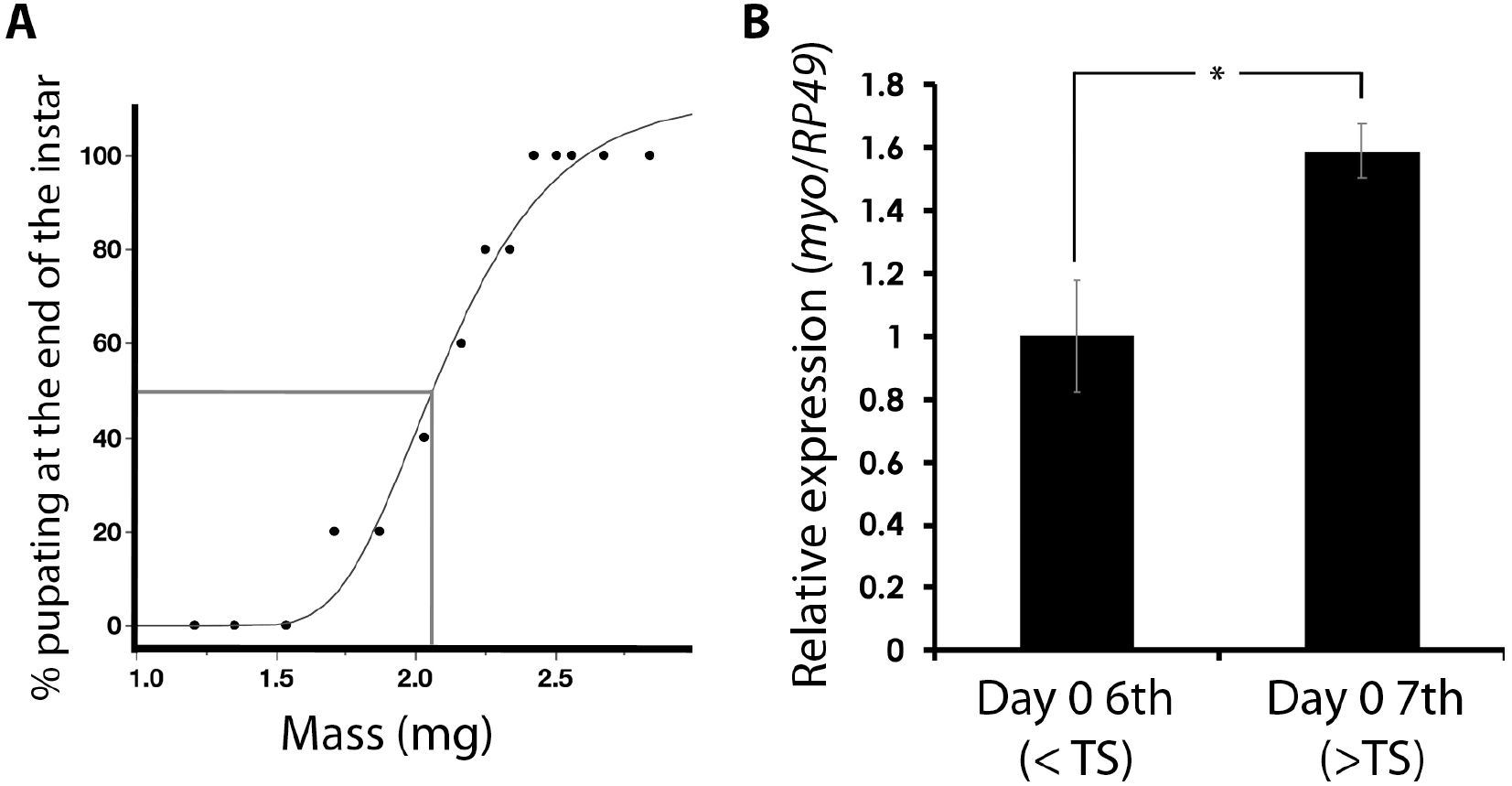
*myo* is upregulated in post-threshold size *T. castaneum* larvae. (A) Determination of threshold size. *T. castaneum* undergo variable number of molts passing through at least seven instars. On day 1 of the final instar, *T. castaneum* larvae undergo pupal commitment (Kamsoi and Belles, 2019). If larvae undergo pupation at the end of the instar, the larval weight on day 1 must be the threshold size. The moving average masses of five day 1 seventh instar larvae were plotted against the percentage of larvae that pupated. Our data show that at 2.1 mg, 50% of larvae have reached the pupal commitment point, indicating that if seventh instar larvae have reached 2.1 mg at the time of the molt, they must have reached the threshold size. (B) Determination of *myo* in freshly molted sixth instar (below the threshold size) and seventh instar larvae above 2.1 mg (above the threshold size). Expression represents the whole body minus the gut and fat body. * denotes statistically significant difference (Student’s t-test: p<0.05).

The *myo* homolog was identified in the *Tribolium* genome and clusters with other Myo homologs in other insects (Fig. S4, S5). We examined the expression of *myo* in freshly molted sixth instar (below the threshold size) or freshly molted final instar larvae weighing at least 2.2 mg (above the threshold size). To do this, we removed the gut and the fat body and examined the expression in the rest of the body (containing the CNS, muscles and integuments). We found a small but significant increase in *myo* expression in the seventh instar larvae (Fig. 6B; Student’s t-test: t(6)=3.02, p<0.05).

To functionally characterize the role of *myo*, we injected *amp^r^* and *myo* dsRNA into larvae. Knockdown verification experiments demonstrated that *myo* dsRNA successfully knocked down the expression of *myo* (Fig. 7A). Eleven out of 18 *amp^r^* RNAi larvae injected in the sixth instar pupated after the seventh instar while four larvae pupated after the eighth instar (Table 1). *amp^r^* dsRNA-injected seventh instar larvae all underwent pupation without a larval-larval molt (n=7; Table 1). Knockdown of *myo* did not induce visible morphological changes. However, larvae injected with *myo* dsRNA as sixth instars continued to molt indefinitely, and those injected as seventh instars never molted; none of these larvae ever entered the prepupal stage (n=18; Table 1). In addition, the intermolt period was significantly increased (Fig. 7B). Overall, the larval duration of *amp^r^* dsRNA-injected larvae was 12 days whereas *myo* dsRNA-injected larvae stayed at the larval stage for up to 7 months before dying (Fig. 7C). When a subset of the larvae was weighed and their fates were assessed, we found that *myo* dsRNA-injected larvae grew at a slower rate than the *amp^r^* dsRNA-injected larvae (Fig. 7D). However, the *myo* dsRNA-injected larvae continued to molt as supernumerary larvae even when they weighed more than the mass of irreversible pupal commitment (i.e. 2.1 mg) one day after the molt and should have pupated (Fig. 7E). In contrast, *amp^r^* dsRNA-injected larvae weighing more than 2.1 mg one day after the molt pupated at the end of the instar (Fig. 7E). These observations indicate that *myo* dsRNA-injected larvae continued to undergo supernumerary molts even after reaching the size when larvae are normally pupally committed, thus supporting the idea that Myo is the signal that mediates the switch between growth and metamorphosis.

**Figure 7.**
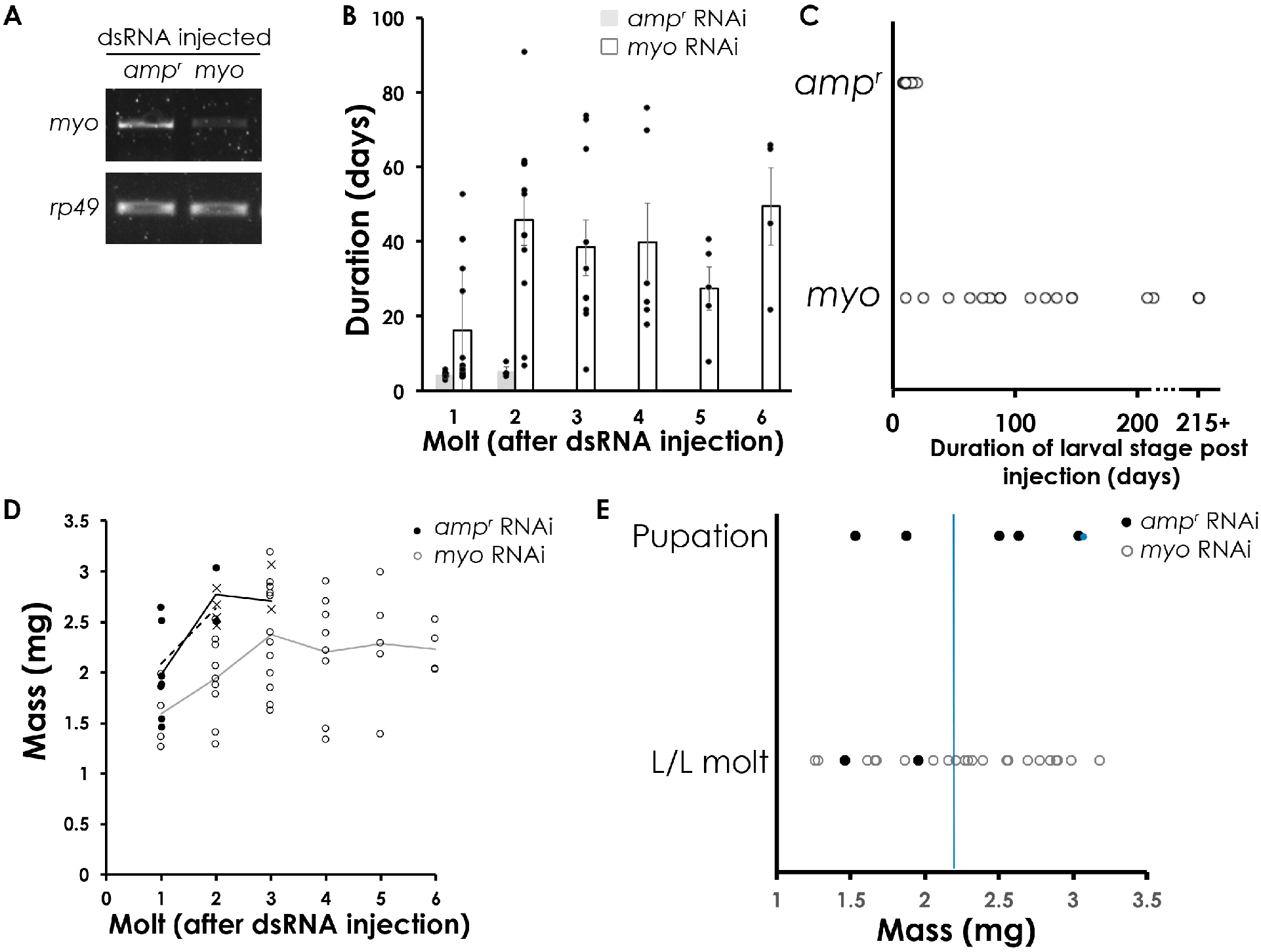
Knockdown of *myo* results in molting delays and prolonged larval stage of *T. castaneum*. **(A)** Knockdown verification showing that *myo* dsRNA injection leads to reduced expression of *myo* in *myo* dsRNA-injected larvae relative to *amp^r^* dsRNA-injected larvae. Cycle number used in gel image: *myo* = 35 cycles; *rp49* = 25 cycles. (B) Duration of each larval instar post dsRNA injection. Bars represent the average duration of each larval instar. Averages were combined for larvae that pupated after the seventh and eighth instar because the duration of each instar did not differ between the groups. *amp^r^* dsRNA-injected larvae were used as the control. Error bars represent standard error. (C) Duration of larval period in *amp^r^* and *myo* dsRNA-injected larvae. Three of the *myo* dsRNA-injected larvae were still alive after 215 days (7 months). Day 0 sixth instar larvae were injected with 0.25*μ*L of dsRNA using a 10*μ*L glass capillary needle. (D) Growth trajectory of *myo* and *amp^r^* dsRNA-injected larvae. Lines represents average larval masses for each time point: black lines represent *amp^r^* dsRNA-injected larvae that underwent two larval molts post dsRNA injections prior to metamorphosis; dotted lines represent *amp^r^* dsRNA-injected larvae that underwent one larval molt post dsRNA injections prior to metamorphosis; gray lines represent *vvl* dsRNA-injected larvae. Filled circles represent *amp^r^* dsRNA-injected larvae; “X” represents prepupal masses of *amp^r^* dsRNA-injected larvae; open circles represent *myo* dsRNA-injected larvae. (E) Fate of larvae at different masses. Larvae were weighed one day after a molt. Filled circles represent *amp^r^* dsRNA-injected larvae; open circles represent *myo* dsRNA-injected larvae. Each larva may be represented by several circles if they continued to undergo multiple supernumerary molts.

**Table 1.** Summary of molting events in *myo* knockdown larvae. Sixth and seventh instar larvae were injected with *myo* or *amp^r^* dsRNA. Injected animals were checked daily for molts, prepupal formation and death.

|  |  |  | Pupating larvae |  |  | Larvae that never pupated<br>and died as larvae |  |  |
| --- | --- | --- | --- | --- | --- | --- | --- | --- |
| dsRNA | Stage<br>injected | #<br>injected | # pupating<br>without<br>molting | # molted<br>once<br>before<br>pupation | # molted<br>twice<br>before<br>pupation | # never<br>molted | # molted<br>once | # molted<br>more<br>than<br>twice |
| *amp<sup>r</sup>* | Day 0 6<sup>th</sup> | 18 | - | 11 | 4 | - | 3 | - |
|  | Day 0 7<sup>th</sup> | 7 | 7 | - | - |  |  | - |
| *myo* | Day 0 6<sup>th</sup> | 18 | - | - | - | 3 | 4 | 11 |
|  | Day 0 7<sup>th</sup> | 6 | - | - | - |  | 4 | 2 |

## DISCUSSION

In this study, we investigated the mechanism by which insects sense their body size and initiate the switch from larval growth to metamorphosis. We found that hypoxia-treatment and a nutrient-deficient diet can both alter threshold sizes in *M. sexta*. The muscle/integument mass was found to be correlated with threshold size: although the relative size of muscles/integuments was greater in hypoxia-treated larvae than that in nutrient-deprived larvae, the same absolute muscle/integument mass was observed at their respective threshold sizes. We found that *myo* expression increases significantly in the muscles during development. Moreover, in the muscles, the expression of *myo* was significantly higher in post-threshold size larvae than in pre-threshold size larvae under both hypoxia and 40% nutrient treatments. *myo* RNAi knockdown in *T. castaneum*, led to permanent indefinite supernumerary larval-larval molts even in larvae that were larger than the threshold size. Based on these findings, we propose that Myo is the signal by which larvae can assess their body size.

### A nutrient-dependent pathway regulates the threshold size

Because the threshold size sets the number of larval instars, the size at metamorphosis and the size of the adult, the threshold size is arguably the most important determinant of final body size. In this study, we found that prolonged exposure to reduced nutrient-diet increases the threshold size (Fig. 1). When larvae were nutritionally deprived during the third instar, the threshold size ranged between 0.65 g to 0.75g. However, when larvae were fed a 40% diet throughout much of the larval instar, the threshold size increased to 0.85 g. Poor nutrient conditions throughout much of the growth period, or recovery from starvation during the third instar, therefore increase the threshold size, suggesting that a nutrient-dependent process likely contributes to the determination of threshold size. In contrast, hypoxia treatment does not reduce the threshold size as larvae reared in hypoxic conditions in either the third or the fourth instar had a threshold size of approximately 0.65 g, similar to larvae that had been fed a 40% diet during the third instar. These findings indicate that 0.65 g may be the “actual” threshold size under normal nutrient conditions.

We discovered that the mass of the muscles/integuments is correlated with threshold size in both hypoxia-treated larvae and nutrition-deprived larvae. The relative muscle/integument mass at threshold size of hypoxia-treated larvae is greater than the relative muscle/integument mass in nutrition-deprived larvae, which have a larger threshold size. Furthermore, the muscle/integument masses at the threshold size of hypoxia-treated and nutrient-deprived larvae are similar. Such correlations are not observed in the fat body mass. These observations indicate that muscle/integument mass serves as a good proxy for body size and the threshold size. Since only the muscle showed consistent increase of *myo* expression in post-threshold size larvae, we think that the expression of *myo* in the muscle is the key signal by which insects assess their body size.

### *Myo* mediates the transition between growth and metamorphosis

Using *T. castaneum*, we found that *myo* RNAi leads to indefinite larval-larval molts. The *myo* dsRNA-injected larvae grew slower compared to the *amp^r^* dsRNA-injected larvae. However, these larvae continued to molt as larvae even when they had reached a mass that normally would have initiated prepupal development. Thus, these larvae clearly had reached the threshold size but were unable to molt into the final instar.

Based on our findings, we propose that Myo couples the attainment of threshold size to the initiation of metamorphosis (Fig. 8). We found that *myo* expression in the muscles increases during the growth phase and is consistently expressed at significantly higher levels in muscles of post-threshold size larvae. The muscle size is a reliable proxy for body size, and the correlated increase in *myo* serves as a signal of body size. Thus, we propose that Myo levels in muscles provide a molecular readout of body size. Upon the attainment of a threshold size, the muscles produce enough Myo to signal to the neuroendocrine center to switch from the growth phase to initiate the physiological processes of metamorphosis. At this point, we do not yet know if Myo levels in the hemolymph stimulate the neuroendocrine glands directly or via another relay system like the nervous system. For example, Myo has been shown to inhibit neuromuscular junction development and synaptic transmission in *D. melanogaster* (Augustin et al., 2017). Thus, it is possible that the signals are transmitted neuronally to the brain or to the corpora cardiaca/corpora allata.

**Figure 8.**
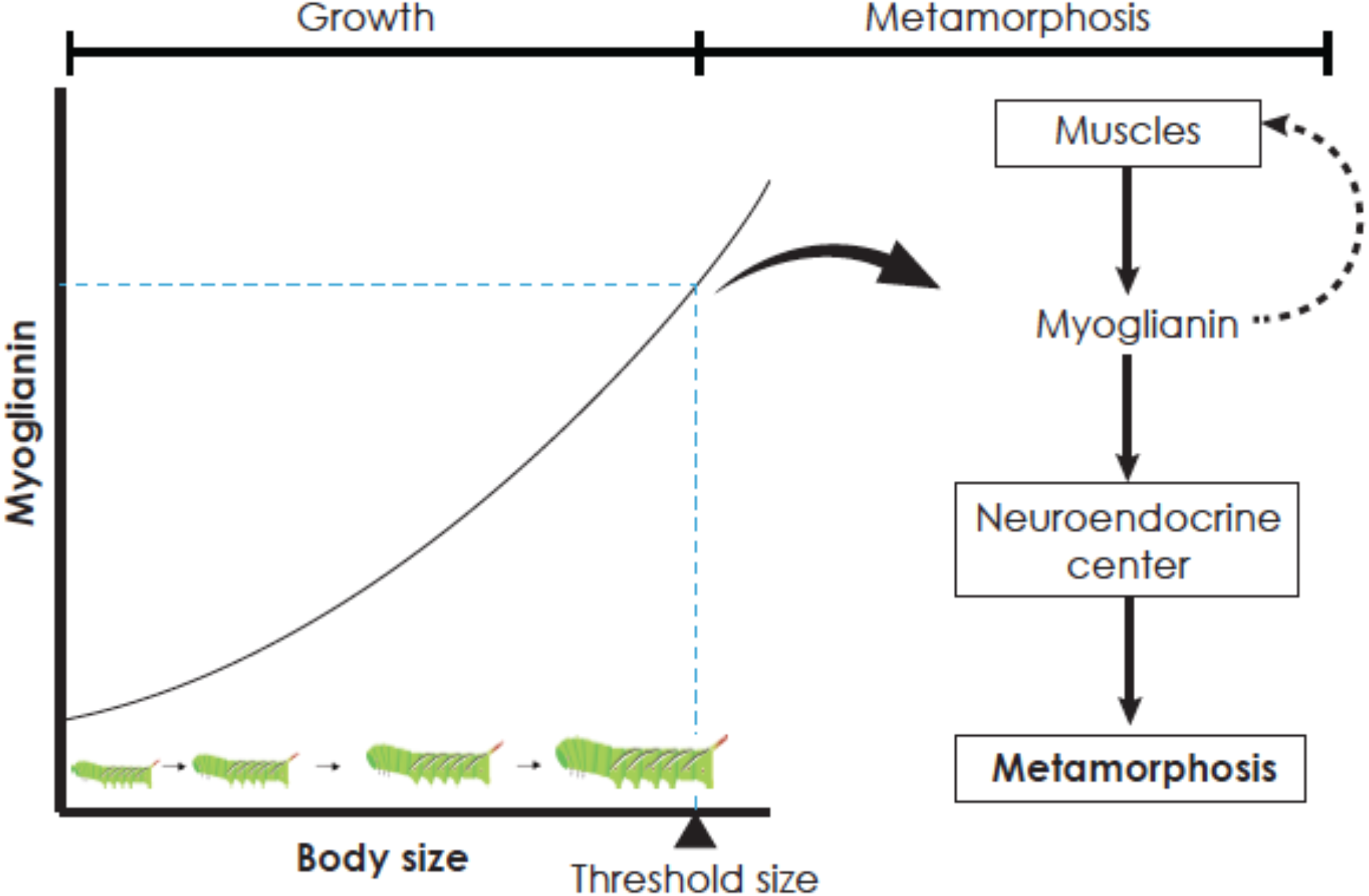
Hypothetical model for how growth of the body is coupled to initiation of metamorphosis. In organisms undergoing determinate growth, the growth phase and the reproductive phases are temporally separated. In this model, the growing tissues (primarily muscles) produce increasing amounts of Myo. Once a threshold level is reached, Myo triggers the end of the growth phase by affecting the neuroendocrine regulators of metamorphosis. Because the same factor acts as both an indicator of growth and initiator of metamorphosis, metamorphosis can be triggered precisely at the threshold size. Dotted line indicates potential positive autocrine feedback.

Taken together, our study demonstrates that one single signaling molecule couples growth to the initiation of metamorphosis and ensures that metamorphosis is triggered when a larva passes the threshold size. Such a model would explain the sharp and precise transition in fates of larvae at the threshold size.

*myo* RNAi has also been conducted in hemimetabolous insects: In both the cricket, *G. bimaculaus*, and the cockroach, *B. germanica*, silencing *myo* leads to multiple supernumerary nymphal molts, often leading to larger body masses (Ishimaru et al., 2016; Kamsoi and Belles, 2019). Thus, in both hemimetabolous and holometabolous insects, *myo* may act as the switch that mediates allows juveniles to shift from the growth phase to the reproductive phase once the animal has reached a specific size threshold. It will therefore be of interest to determine whether a threshold size can be identified in hemimetabolous insects.

Finally, JH-deficient larvae of *T. castaneum* and *B. mori* must undergo at least three larval molts to produce an unknown “competence factor” that allows them to initiate metamorphosis (Daimon et al., 2015; Smykal et al., 2014). Whether or not Myo is related to the “competence factor” is unclear at this point.

### JH does not affect threshold size of *M. sexta*

When we applied JH during the fourth instar of *M. sexta*, we observed that the threshold size could not be shifted (Fig. S6). Thus, we believe that JH is not the primary regulator of threshold size in *M. sexta*. Instead, JH is a downstream effector that mediates the decision to metamorphose post-threshold size attainment.

A previous study has suggested that threshold size in *T. castaneum* might be regulated by JH (Chafino et al., 2019). However, in that study, the threshold size was identified by starving larvae at various sizes during the final instar and identifying the size above which the larva initiates metamorphosis. Whether this size checkpoint is equivalent to the threshold size checkpoint remains unclear. While the clearance of JH is necessary to initiate metamorphosis, the threshold size assessment must occur earlier. Myo has been shown to act upstream of JH signaling in hemimetabolous insects: *myo* knockdown leads to an increase in *jhamt* expression in the corpora cardiaca/corpora allata (Kamsoi and Belles, 2019). Thus, Myo is likely the threshold size determinant that ultimately causes a drop in JH titer.

*T. castaneum* and *M. sexta* differ in their response to nutrient manipulations in younger larvae. In *T. castaneum*, feeding larvae on flour that has been diluted to 20% during the fifth instar leads to a lack of molt and eventual metamorphosis, similar to a bail-out response seen in other beetles (Nagamine et al., 2016; Shafiei et al., 2001; Terao et al., 2015). *M. sexta* does not appear to exhibit this type of bail-out response as they either molt or die when nutrients are removed. The bail-out response seen in *T. castaneum* likely represents an adaptive response that is absent in *M. sexta*.

### Conserved functions of Myo in *T. castaneum* and other insects

In addition to the role of Myo during the threshold size checkpoint, *myo* knockdown in *T. castaneum* revealed additional functions of Myo that are conserved across insects. When *myo* was knocked down in *T. castaneum*, the intermolt period was dramatically increased, with some larvae taking a month to molt. Normally, the intermolt period is around five days. Thus, *myo* knockdown delays the onset of a molt, presumably through a decrease in ecdysteroidogenesis. Activin signaling has previously been shown to affect ecdysteroidogenesis in both *B. germanica* and *D. melanogaster* (Gibbens et al., 2011; Santos et al., 2016). Thus, the role of Activin signaling on ecdysteroidogenesis appears to be conserved across various insect species.

In addition, we observed that when Myo was silenced, the larva grew slower, indicating that it promotes growth. This growth promoting function of Myo appears to be conserved in most insects (Ishimaru et al., 2016; Kamsoi and Belles, 2019). The only exception is found in *D. melanogaster*, where an opposite effect is observed: In *D. melanogaster*, knockdown of *myo* in the muscles leads to a larger larval size whereas overexpression of *myo* in the muscles leads to smaller larval body size without affecting the developmental time (Augustin et al., 2017). In this species, the functions of TGF-beta ligands appear to be switched with *Activin-beta* playing a growth promoting role, similar to the growth-promoting role of *myo* in other insect species: the loss-of-function mutation in the *Activin-beta* gene leads to smaller muscle and adult body size in *Drosophila* without affecting the timing of metamorphosis (Moss-Taylor et al., 2019). Thus, the growth promoting roles of TGF-beta ligands may have been switched in the lineage leading to *D. melanogaster.* Such switches in function have also been reported for other TGF-beta ligands (Namigai and Suzuki, 2012).

### Implications for mammalian growth

Body size regulation in insects also has many parallels to size regulation in mammals. Puberty in mammals, like metamorphosis in insects, is initiated upon reaching a specific threshold size, and its onset is affected by body size and nutritional status (Hirsch and Batchelor, 1976). Obesity in children leads to precocious pubertal onset, a public health issue that is affecting US youths (Burt Solorzano and McCartney, 2010; Euling et al., 2008). While being overweight can lead to precocious puberty, we do not know if the threshold size itself is shifted under altered nutritional regimes. Based on our study, we hypothesize that the precocious pubertal onset may be a product not only of precocious attainment of body size but potentially also of an adjustment of the threshold size itself. Whether the vertebrate homologs of Myo, BMP-11/GDF-8, mediate the transition between pre-pubertal growth and puberty has not been clearly demonstrated although a polymorphism in GDF-8 has recently been shown to delay puberty in cows (Cushman et al., 2015). Additional studies are necessary to demonstrate whether an evolutionarily conserved mechanism is involved in the regulation of determinate growth.

## CONCLUSION

In this study, we sought to identify the mechanism by which larvae assess their body size. Specifically, we investigated mechanism by which larvae sense the threshold size, the first size check point that determines the timing of metamorphosis. Muscle growth and the expression of the *myo* in the muscles are linked to the attainment of the threshold size, the earliest body size checkpoint for metamorphosis.

## METHODS

### Animal rearing

Wildtype *M. sexta* were obtained from Carolina Biological Supply Company. Larvae were fed a standard artificial diet as described previously (Kemirembe et al., 2012) unless otherwise noted. Larvae were raised in individual plastic cups at 26.5°C and a 16:8 hr light:dark cycle. Larvae were kept in 1 oz soufflé cups until the end of the fourth instar, when they were moved to a 5 oz soufflé cup. The end of an instar can be clearly identified in this species when the head capsule begins to slip (denoted “HCS” for head capsule slippage) as the larva initiates a molt. Wildtype GA-1 strain *Tribolium castaneum* beetles were raised on organic whole wheat flour supplemented with 5% nutritional yeast and 0.5% fumagilin at 29.5°C and ~55% humidity.

### Hypoxia treatment

*M. sexta* larvae subjected to hypoxia treatments were moved to an airtight cell culture chamber immediately after molting into either the third or the fourth instar and kept in cups with multiple holes in the lid. A 5% oxygen/carbon dioxide mixture was sent into the chamber and oxygen levels were kept at approximately 4±1% throughout the experiment. Larvae were removed from the chamber at the end of the instar, after which the artificial diet was replaced with fresh diet, and multi-holed lids were exchanged for single-holed lids.

### Effects of nutrients on threshold size

To assess the effects of nutrition on growth and threshold size, larvae were removed from the standard artificial diet and placed onto experimental diets or subjected to starvation conditions at the onset of the third or fourth instar. The experimental diet contained 40% the amount of dietary protein compared to the standard diet by having reduced levels of wheat germ and casein (Table S1). Larvae subjected to starvation conditions were removed from the standard diet 24 hrs after molting into the third instar and placed onto a moistened Kimwipe, which served as a water source. All larvae were returned to the standard artificial diet when they initiated HCS.

### Methoprene treatment

To test the effect of JH on threshold size, third instar larvae were hypoxia-treated at the onset of the third instar as described above. Two days after the onset of third instar HCS, 1 *μ*l of methoprene (Sigma) dissolved in acetone (10 *μ*g/*μ*l) was applied topically on the dorsal side of the fourth instar. These larvae were weighed at fourth instar HCS and subsequently tracked to see if they entered the wandering stage – an early indication of the onset of metamorphosis – or initiated another molt to determine the threshold size.

### Threshold size determination

To determine how varying growth conditions altered the threshold size, *M. sexta* larvae were subjected to varying nutritional or hypoxic conditions, and their growth was followed. Subjecting larvae to these sub-optimal growth conditions generates larvae above and below threshold size at the end of the fourth instar, and threshold size can be determined by plotting their developmental fate against their mass at that time. Larvae were moved onto experimental diets at the onset of the third or fourth instar or subjected to starvation or hypoxic conditions as described above. Larvae were checked daily, starting on the day of HCS into the fourth or fifth instar, for third or fourth instar hypoxia-treated larvae, respectively. Observations continued until larvae initiated HCS into a supernumerary instar or exhibited signs of metamorphosis, as indicated by a purging of the gut contents and development of a darkened dorsal vessel. The percentage of larvae that entered final larval instars was plotted against mass at the time of the fourth HCS and fitted to a sigmoidal growth curve. For each treatment, the mass at which 50% of the larvae entered the final instar was determined to be the threshold size.

### Muscle and fat body mass

After the larvae were weighed, they were dissected in 1X phosphate-buffered saline (PBS; 0.15 M NaCl, 0.0038 M NaH_2_PO_4_, 0.0162 M Na_2_HPO_4_; pH 7.4). The gut, central nervous system (CNS) and other tubular structures were removed. The fat body was then carefully removed and placed on an aluminum foil, dried at 60°C for at least 48 hrs, and then weighed. The rest of the larval body, representing the combination of muscles and integuments, was also dried and weighed.

### RNA isolation and cDNA synthesis

To determine the expression of *myo* at the end of various instars, RNA was isolated from the anterior half of the larva (including the head and the thorax) at the onset of HCS when the head capsule was still fluid filled and mandibles were white; three biological replicates were created for each time point. To determine the expression of *myo* in pre-threshold size and post-threshold size larvae, larvae were either placed on 40% diet once they molted into the second instar or placed in hypoxia conditions for the duration of the third instar. Muscles, fat body and CNS were dissected from *Manduca* at the onset of HCS in the fourth instar in 1X PBS. To isolate RNA, tissues were homogenized in 500 *μ*L of TRIzol reagent (Thermo Fisher). After extracting RNA using chloroform, the RNA was treated with DNAse (RQ1 RNase-Free DNase, Promega) to remove remaining traces of genomic DNA. The First Strand cDNA Synthesis Kit (Thermo Fisher) was used to convert 1 *μ*g of RNA to cDNA via reverse transcription.

### Quantitative polymerase chain reaction (qPCR)

SYBR Green Supermix (Bio-Rad) and qPCR primers (Table 2) were used for qPCR. To measure *myo* expression, primers targeting the *Manduca* homolog of the *Drosophila myo* gene (Genbank accession no. XM_030169402.1), were designed. Each replicate was assayed in triplicate with no-template controls. *RpL17A* was used as an internal control gene, and a standard curve method was used to analyze the data. JMP (SAS Institute, Cary, NC) was used for statistical analyses.

### Double-stranded RNA (dsRNA) synthesis

*T. castaneum myo* gene (Genbank accession nr: XM_961726.3) was amplified using the primer set listed in Supplemental Table 1. The PCR product was inserted into a pCR4 TOPO vector (Thermo Fisher Scientific) following the manufacturer’s instructions. Plasmid DNA from transformed *E. coli* cells were isolated according to the QIAprep Spin Miniprep Kit protocol (Qiagen). A restriction reaction was then set up to linearize the plasmid DNA. Using 1 *μ*g of linearized plasmid as a template, ssRNA was then synthesized using the MEGAscript T3 and T7 kits. The synthesized ssRNA was cleaned using phenol/chloroform extraction and annealed to make a 2 *μ*g/*μ*l solution as described previously (Hughes and Kaufman, 2000). Proper annealing was checked using gel electrophoresis.

### Microinjection of *T. castaneum* larvae

Before injection with dsRNA, sixth and seventh instar *T. castaneum* larvae were first anesthetized on ice. A 10*μ*L glass capillary needle was used to inject 0.5*μ*L of dsRNA into seventh instar larvae and 0.25*μ*L into sixth instar larvae. Control animals were also injected with the same volume of bacterial *ampicillin-resistance (amp^r^)* dsRNA.

### Knockdown verification

In order to verify proper knockdown of *myo*, day 0 sixth instar *T. castaneum* larvae were injected with either *myo* or *amp^r^* dsRNA. RNA from three day 4 sixth instar larvae were collected and processed as outlined above. After converting 1 *μ*g of RNA into cDNA, PCR was run using *myo* and *rp49* primers listed in Table 2. For *rp49*, 20, 25 and 30 cycles were run; for *myo*, 30, 35 and 40 cycles were run.

## Acknowledgements

We thank Drs. Kimberly O’Donnell, Louise Darling, Melissa Beers and Julie Roden for their helpful advice and discussions. We also thank the members of the Suzuki lab and Heidi Park for their support and comments on the manuscript.

## SUPPORTING INFORMATION

**Supplemental figure S1.**
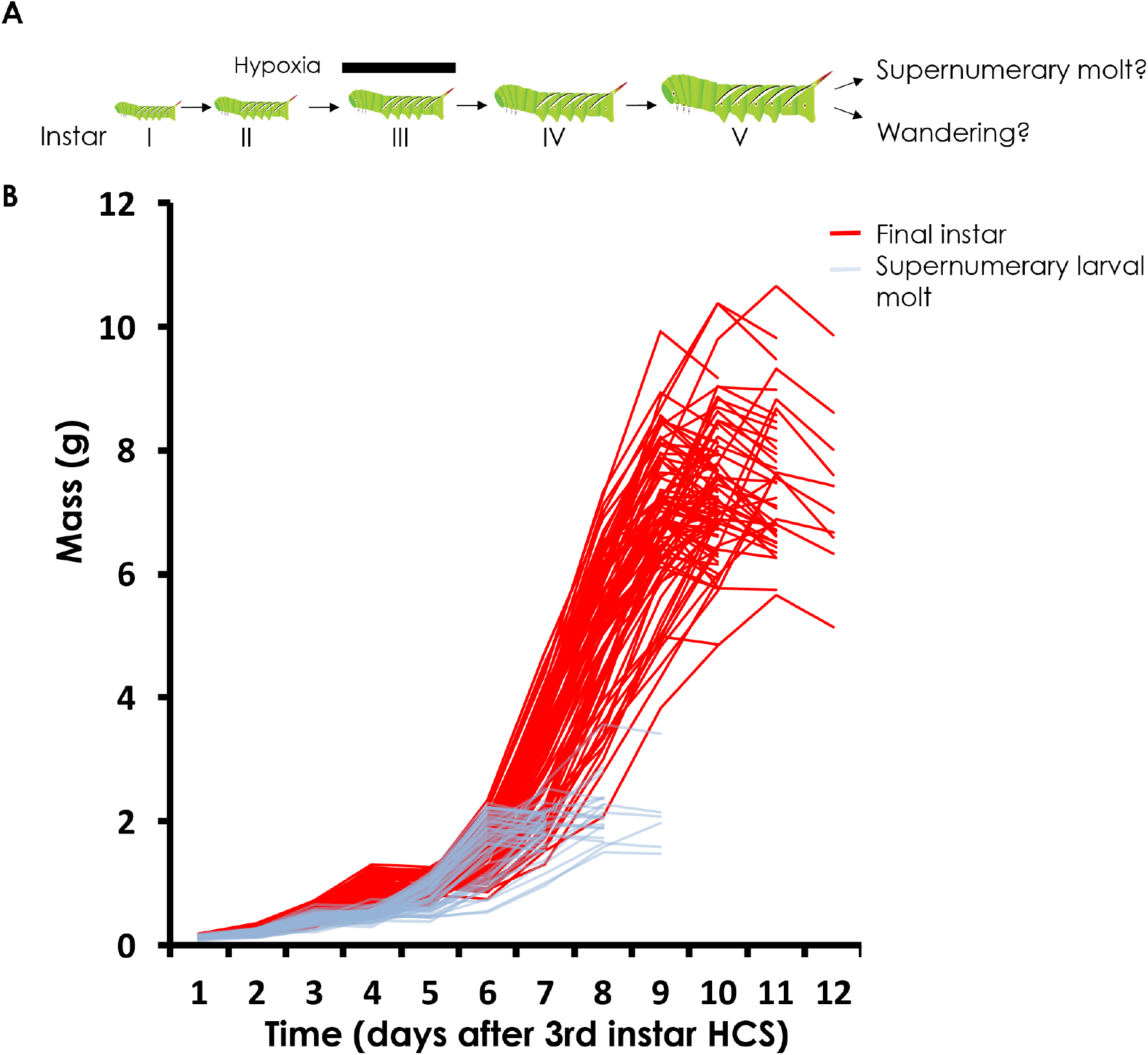
Hypoxia generates two developmental trajectories for *Manduca sexta*. (A) A scheme showing the timing of hypoxia treatment. (B) Individual growth trajectories of larvae that underwent wandering (red) or supernumerary molt at the end of the fifth instar (light blue). Larvae were subjected to hypoxic conditions from the beginning to the end of the third instar. Masses were recorded at the end of the third instar and every day after.

**Supplemental figure S2.**
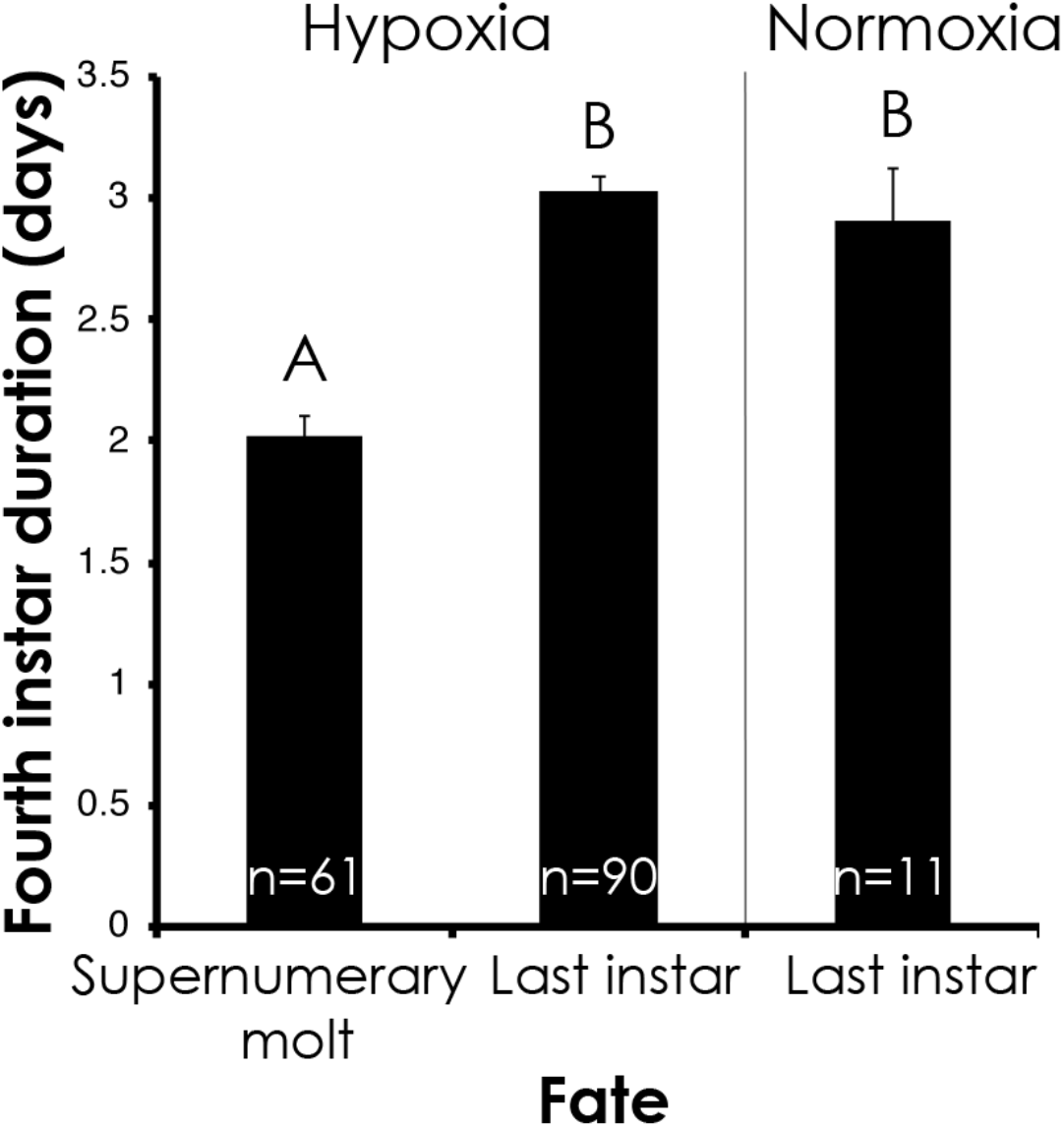
Fourth instar duration predicts developmental fate of the larva. Fourth instar duration (mean±SE) of larvae that wandered at the end of the fifth instar (denoted “last instar” and those that underwent a supernumerary molt (denoted “supernumerary larval molt”) are shown for larvae that were reared under hypoxic or normoxic conditions during the third instar. Days were counted from day 0 of the fourth instar. One-way ANOVA: F(2,159) = 125.556, p<0.0001. Means not sharing the same letter are statistically significant (Tukey HSD, p<0.0001).

**Supporting figure S3.**
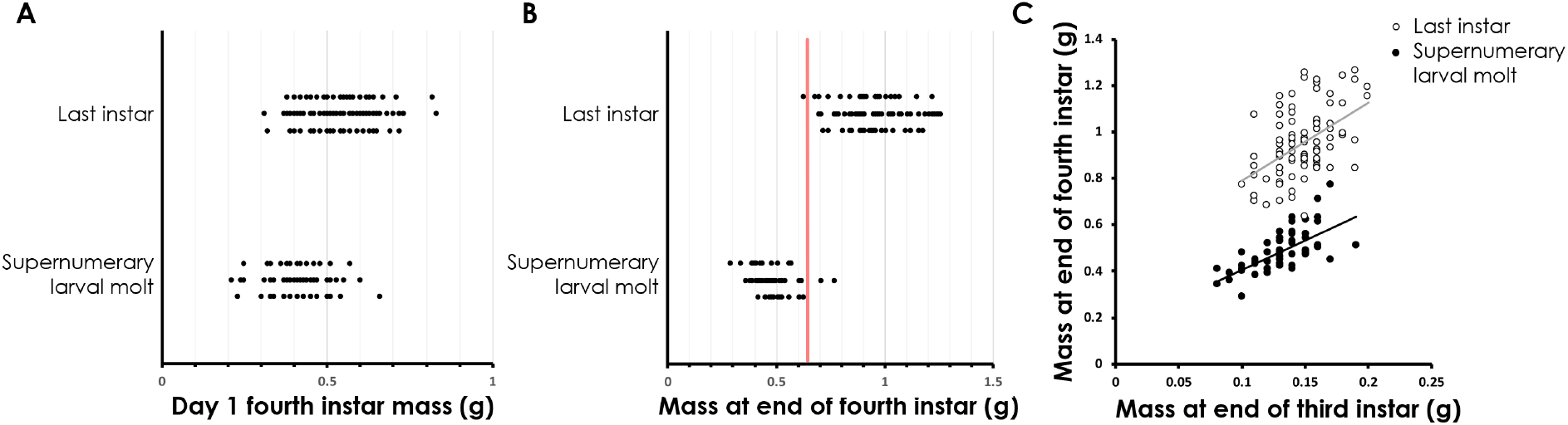
Mass at the end of the fourth instar predicts the developmental fates of larvae. (A) The mass at first day of the fourth instar is a poor predictor of nature of subsequent molt. “Supernumerary larval molt” denotes fifth instar larvae that molted into another larval instar. “Last instar” denotes fifth instar that initiated wandering. (B) The decision to enter supernumerary stage is made at the end of the fourth instar. Red line indicates the estimated threshold size. (C) A plot of mass at the end of the third instar vs mass at the end of the fourth instar, showing that the mass at the end of the third instar is a poor predictor of threshold size. Filled circles are larvae that underwent a supernumerary larval molt; open circles denote larvae that wandered at the end of the fifth instar. All larvae were subjected to hypoxic conditions during the third instar and then tracked for supernumerary molt/wandering.

**Supplemental figure S4.**
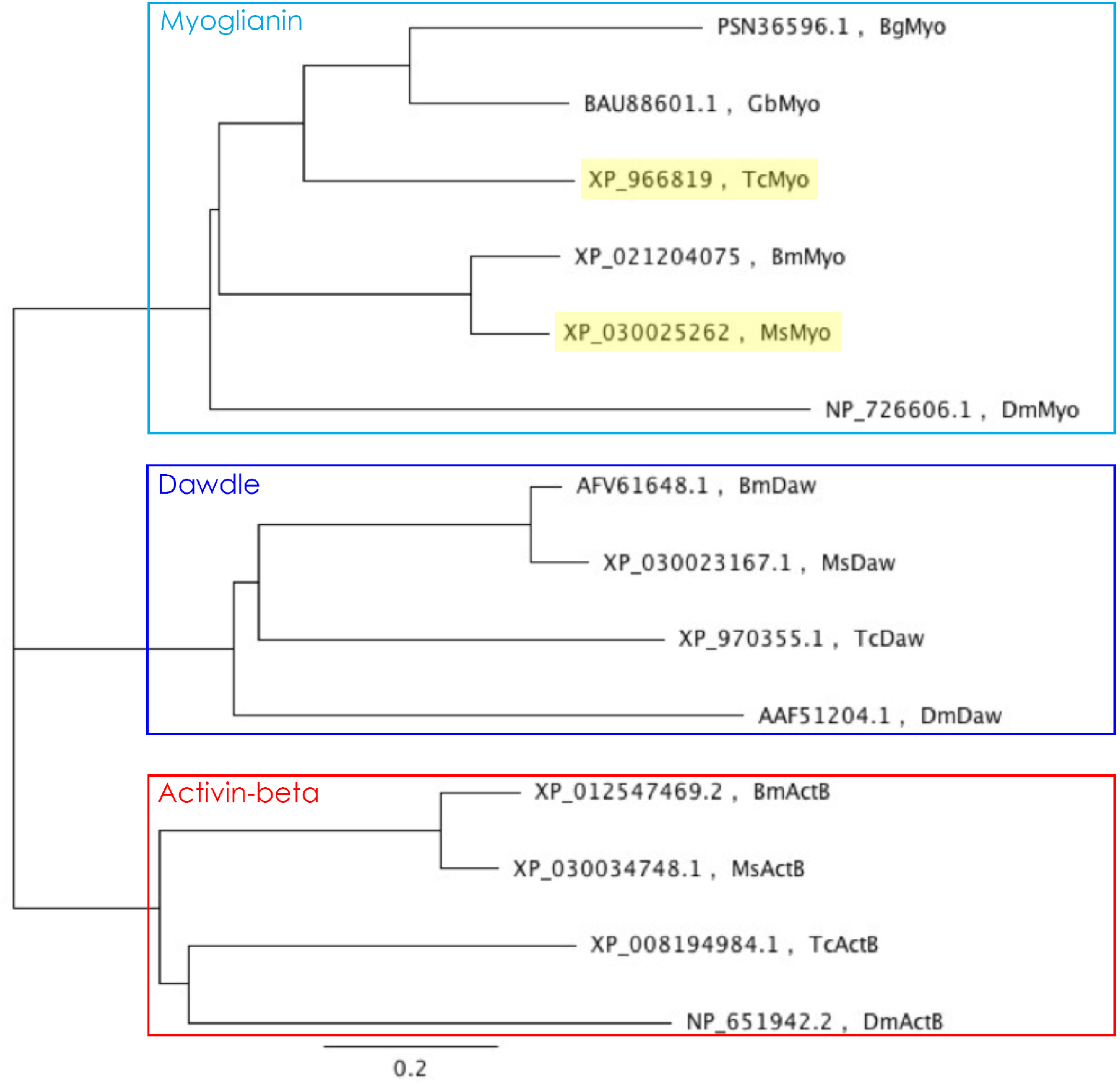
Phylogenetic tree of the three Activin ligands, Activin-beta (ActB), Dawdle (Daw) and Myoglianin (Myo) predicted from genome sequences of *Drosophila melanogaster* (*Dm*), *Blattella germanica* (*Bg*), *Gryllus bimaculatus* (*Gb*), *Tribolium castaenum* (Tc), *Bombyx mori* (*Bm*) and *Manduca sexta* (*Ms*).

**Supplemental figure S5.**
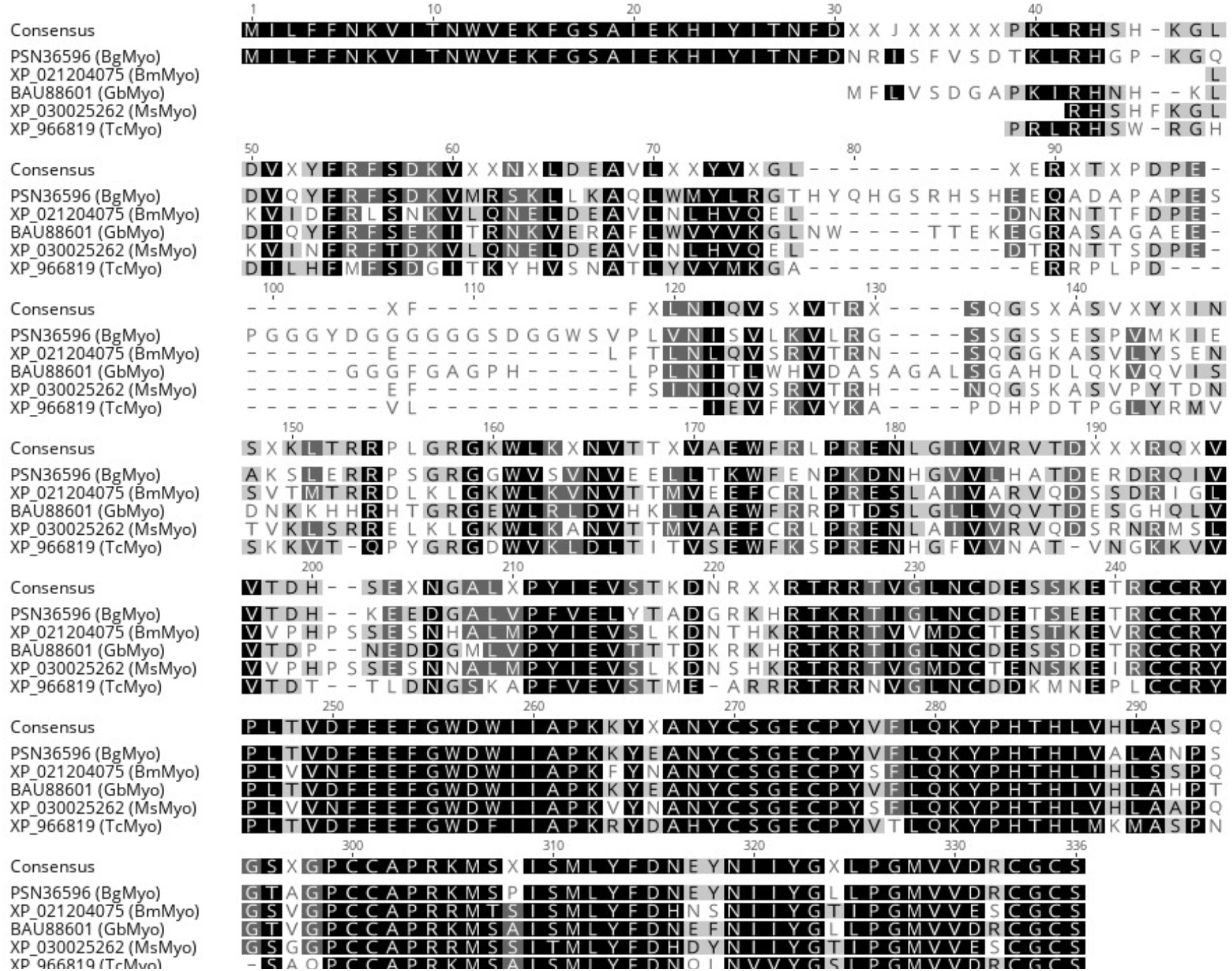
Amino acid alignment of Myoglianin from *Blattella germanica* (*Bg*), *Bombyx mori* (*Bm*), *Gryllus bimaculatus* (*Gb*), *Manduca sexta* (*Ms*) and *Tribolium castaenum* (Tc).

**Supporting figure S6.**
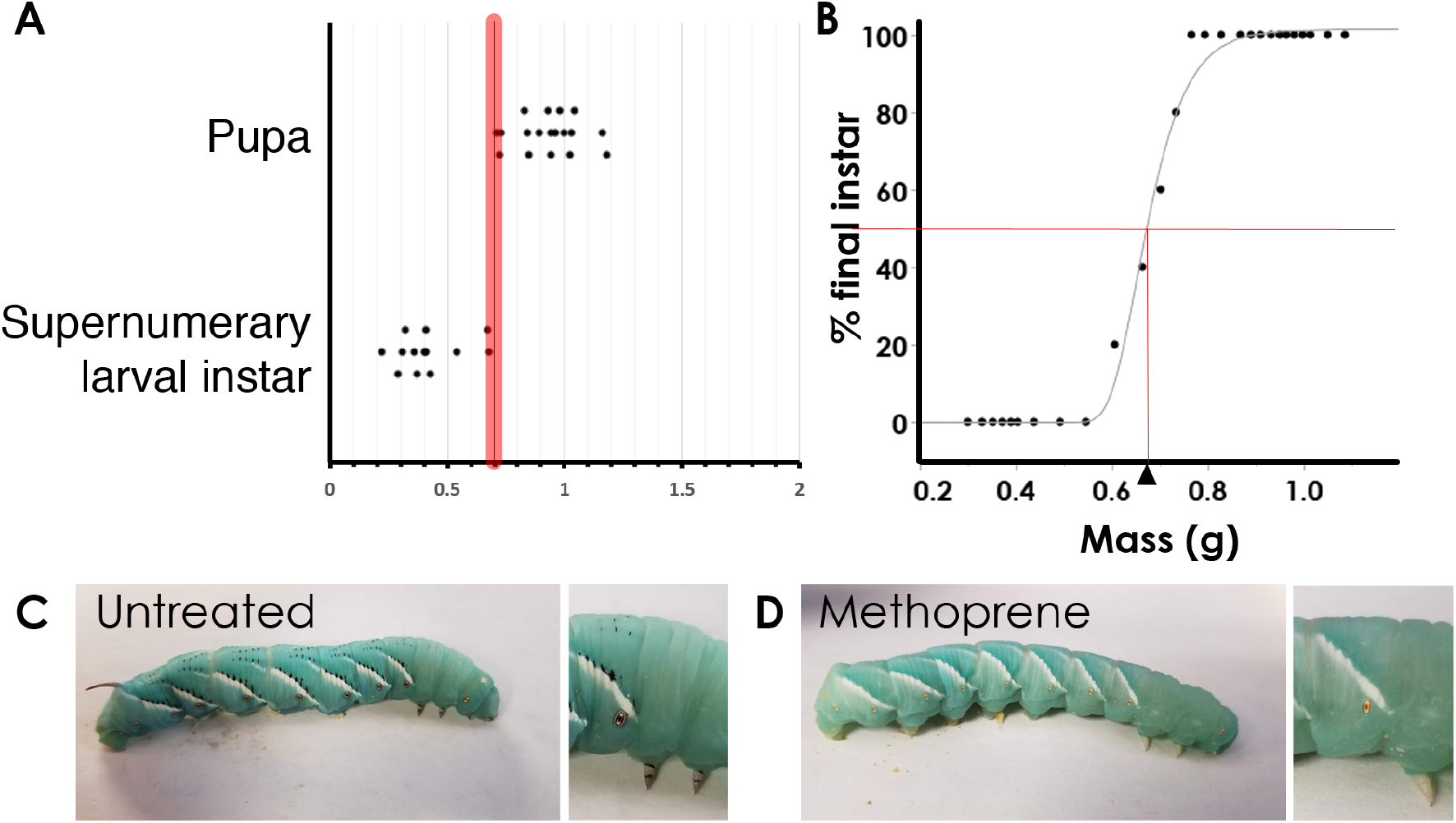
Methoprene does not shift the threshold size. (A) Threshold size determination in methoprene-treated larvae that had undergone hypoxia treatment during the third instar. (B) The average masses of five larvae at the end of the fourth instar were plotted against the percentage of larvae that entered final larval instar. Triangle indicates the threshold size when 50% of the larvae wander at the end of the fifth instar. Line represents a Gompertz 3P model fit. (C) A normal fifth instar larva, showing the melaninic markings on the dorsal side and the legs. (D) A fifth instar that had been treated with methoprene. This larva weighed 1.18 g at 4th HCS, well above the threshold size. Methoprene was applied on the dorsal side two days after 3rd instar HCS.

**Table S1:**
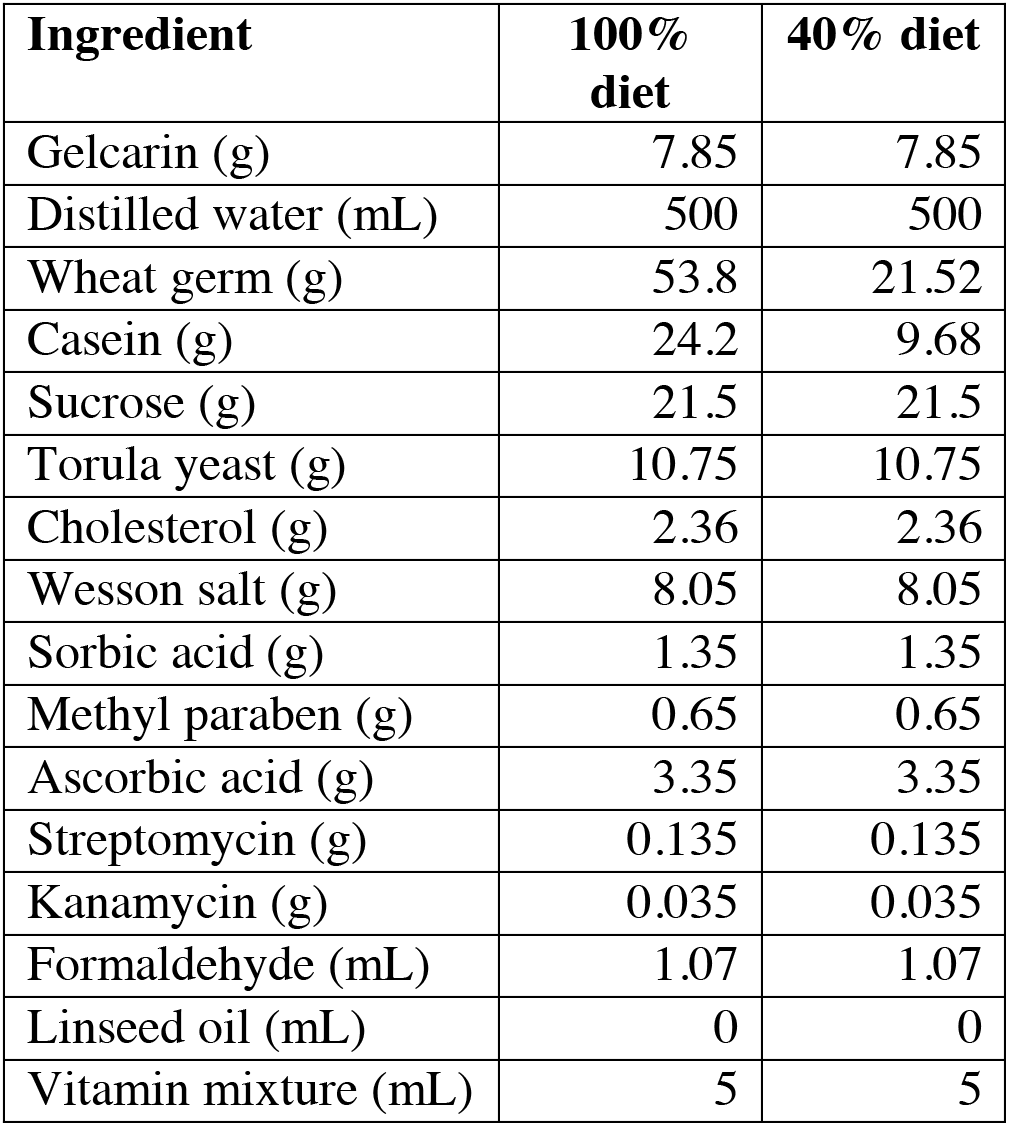
Ingredients for experimental diets used in this study. Percentages based on a standard (100%) diet described by Yamamoto et al (1969)

| Ingredient | 100% diet | 40% diet |
| --- | --- | --- |
| Gelcarin (g) | 7.85 | 7.85 |
| Distilled water (mL) | 500 | 500 |
| Wheat germ (g) | 53.8 | 21.52 |
| Casein (g) | 24.2 | 9.68 |
| Sucrose (g) | 21.5 | 21.5 |
| Torula yeast (g) | 10.75 | 10.75 |
| Cholesterol (g) | 2.36 | 2.36 |
| Wesson salt (g) | 8.05 | 8.05 |
| Sorbic acid (g) | 1.35 | 1.35 |
| Methyl paraben (g) | 0.65 | 0.65 |
| Ascorbic acid (g) | 3.35 | 3.35 |
| Streptomycin (g) | 0.135 | 0.135 |
| Kanamycin (g) | 0.035 | 0.035 |
| Formaldehyde (mL) | 1.07 | 1.07 |
| Linseed oil (mL) | 0 | 0 |
| Vitamin mixture (mL) | 5 | 5 |

**Table S2:**
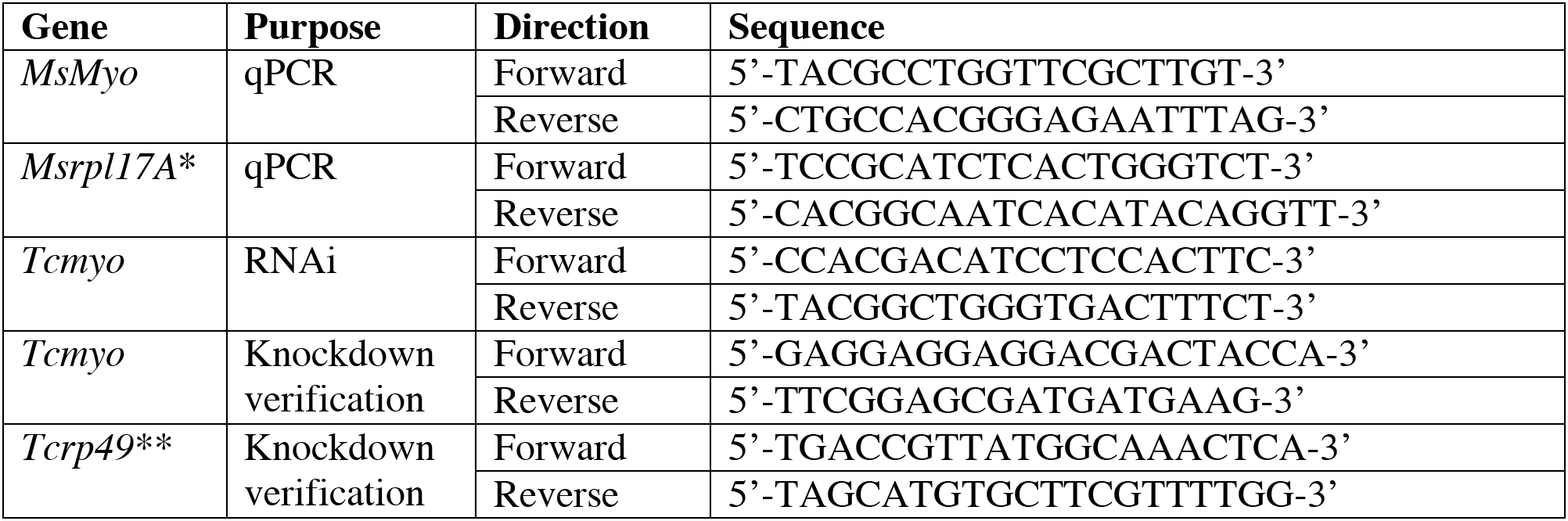
Primer sequences used in this study. *Sequences from Ono et al (2006). ** Sequences from Parthasarathy et al (2008).

